# Genetic architecture of the thermal tolerance landscape of *Drosophila melanogaster*

**DOI:** 10.1101/2024.07.01.601629

**Authors:** Juan Soto, Francisco Pinilla, Patricio Olguín, Luis E. Castañeda

## Abstract

Increased environmental temperatures associated with global warming strongly impact the natural populations of ectothermic species. Therefore, it is crucial to understand the genetic foundations and evolutionary potential of heat tolerance. However, heat tolerance and its genetic components depend on the methodology, making it difficult to predict the adaptive responses to global warming. Here, we measured the knockdown time for 100 lines from the *Drosophila* Genetic Reference Panel (DGRP) at four different static temperatures, and we estimated their thermal death time (TDT) curves, which incorporate the magnitude and the time of exposure to thermal stress. Through quantitative genetic analyses, it was observed that the knockdown time showed a significant heritability at different temperatures and that its genetic correlations decreased as temperatures were dissimilar. Significant genotype-by-sex and genotype-by-environment interactions were noted for knockdown time. We also discovered heritable variation for the two parameters of TDT: *CT_max_* and thermal sensitivity. Taking advantage of the DGRP, we performed a GWAS and identified multiple variants positively associated with the TDT parameters, which map to genes related to signaling and developmental functions. We performed functional validations for some candidate genes using RNAi, which revealed that genes such as *mam*, *KNCQ,* or *robo3* affect the knockdown time at a specific temperature but are not associated with the TDT parameters. Ultimately, the thermal tolerance landscape must possess the ability to adapt to the selective pressures caused by global warming. Genetic variation of phenotypic plasticity and sexually antagonistic pleiotropy should also facilitate the adaptive process and maintain the genetic diversity in *Drosophila* populations.

## INTRODUCTION

Environmental temperature has increased by about 1.1°C during the last century, but global warming effects have been more strong during the last 30 years (Gulev et al., 2021; Hartmann et al., 2013). This scenario is accompanied by an increase in the intensity and frequency of extreme weather events at the local scale, such as frost and heat waves (Meehl & Tebaldi, 2004). These changes exert important selective pressures on natural populations (Pacifici et al., 2015), leading to changes in their distribution (Lenoir et al., 2020) and potentially affecting their persistence over time (Thomas et al., 2004). These changes particularly affect ectotherms, as most already maintain a body temperature close to their thermal limit (Angilletta, 2009; Deutsch et al., 2008). Given this scenario, it is important to understand the genetic architecture of thermal tolerance responsible for heritable phenotypic variability, i.e., the number of genetic variants affecting the trait, their frequencies in the population, the magnitude of their effects, and their interactions with each other and the environment (Timpson et al., 2017). Understanding the genetic architecture of thermal tolerance will make it possible to assess the evolutionary capacity of populations to respond to global warming.

To study thermal tolerance and its genetic determinants, a method is needed that can correctly determine the upper thermal limits of organisms. In this sense, there has been a controversy in recent years as to which is the most appropriate method since numerous studies have observed that the methodology used in the laboratory affects the estimates of the thermal tolerance parameters and their heritability. This makes it challenging difficult to compare findings across studies (Castañeda et al., 2019; Chown et al., 2009; Santos et al., 2012; Sgrò et al., 2010; Terblanche et al., 2007). In general, two methods are used to determine a single parameter to determine the upper thermal limit: (1) static assays, where the organism is exposed to a constant temperature until it collapses (i.e., knockdown time); and (2) dynamic assays, where the organism is exposed to a thermal ramp until it collapses (i.e., knockdown temperature) (Beitinger & Lutterschmidt, 2011). Longer trials increase the effect of other experimental variables, such as nutrient and water availability, during the trial (Mitchell & Hoffmann, 2010; Overgaard et al., 2011). This increases the residual or environmental variability in the assays and, therefore, reduces heritability estimates (Castañeda et al., 2019). With the aim of unifying methodologies in the determination of thermal tolerance, Rezende et al. (2014) proposed using the thermal-death-time (TDT) curves to describe the thermal tolerance landscape of ectotherm species, incorporating both the magnitude and the time of exposure to thermal stress. This integrative approach estimates the critical temperature maximum (*CT_max_*) and thermal sensitivity (*z*) using the knockdown time obtained at different static temperatures. The thermal tolerance landscape provides a unified framework for studying thermal tolerance (Rezende et al., 2020) and links thermal tolerance to cumulative heat injury sustained under natural heat stress (Jørgensen et al., 2019, 2021; Li et al., 2023; Ørsted et al., 2022). However, little is known about the genetic determinants of the TDT curves, which limits our understanding of the evolution of the thermal tolerance landscapes.

Several studies have identified genes associated with a single parameter of thermal upper limits determined by dynamic or static assays (Hoffmann & Willi, 2008). The first genes identified were those encoding heat shock proteins, increasing their expression and reducing the detrimental effects of heat stress (Anderson et al., 2003; Dahlgaard et al., 1998; Lerman & Feder, 2001; McColl & McKechnie, 1999). Other genes also play a role in thermal tolerance. For example, genes involved in longevity and stress response, such as *methuselah* (*mth*) and the transcription factor *foxo* (Araújo et al., 2013; Giannakou et al., 2004; Lin et al., 1998; Morgan & Mackay, 2006), and genes involved in the pathway of hormones and neurotransmitters of the catecholamine family such as *catsup* and *ddc* (Norry et al., 2007, 2009). The role of these genes in thermal tolerance has been validated using field release and recapture experiments (Loeschcke et al., 2011) and gene mutagenesis (Hoffmann & Willi, 2008). In addition, many studies have described extensive lists of genes with variants associated with latitudinal or seasonal variation that may be involved in thermal tolerance (Fabian et al., 2012; Kapun et al., 2020; Kolaczkowski et al., 2011; Machado et al., 2016; Rudman et al., 2022; Zhao et al., 2015). In recent years, the development of genomic tools has facilitated the study of genetic variants associated with phenotypic traits, including the *Drosophila* Genetic Reference Panel or DGRP (MacKay et al., 2012). The DGRP is a set of fully sequenced isogenic lines that facilitates the development of quantitative genetic analysis and genome-wide association analysis, or GWAS, which involves testing the association of genetic variants in the genome of multiple individuals to identify genotypic associations with a given phenotype (Tam et al., 2019). Using this genetic panel, Rolandi et al. (2018) described variants in genes related to cell organization, cell trafficking, and neurotransmitter activity, while Lecheta et al. (2020) classified the genes associated with thermal tolerance into three groups: (1) genes involved in protecting the organism, (2) genes involved in gene regulation of the thermal response, and (3) genes that alter the degree of preparation of the organism’s body to withstand heat stress. While these two studies have used the DGRP to study thermal tolerance using dynamic methods, no work has been done specifically to study the genetic determinants of the parameters of the thermal tolerance landscape.

Identifying the genes and genetic variants that affect thermal tolerance is only part of the genetic architecture; how this genetic determinant interacts with the environment is critical to understanding how organisms respond to thermal variation. Different genotypes can differ in their phenotypes in response to environmental variation, known as genotype-by-environment interaction (GxE) (Saltz et al., 2018), and the presence of this interaction allows for the evolution of different phenotypic optima in different environments (Gillespie & Turelli, 1989). Numerous studies have shown that thermal tolerance to low and high temperatures shows significant GxE (Delclos et al., 2021). Whereas the genetic determinants of thermal tolerance are not fixed but depend on the environment (Ørsted et al., 2019), making GxE a key factor in understanding the evolutionary process under variable thermal environments. Some authors even describe a triple interaction between genotype, environment, and sex in thermal tolerance (Delclos et al., 2021), and that sex is also an important context with which genetic determinants interact (Lasne et al., 2018; Morgan & Mackay, 2006). The existence of a genotype-by-sex interaction (GxS) could imply that there are selective differences between the sexes, with important implications for the evolution and maintenance of genetic diversity in natural populations, as well as the use of resources and habitats in nature (Connallon & Clark, 2013; Lynch & Walsh, 1998; Pennell et al., 2016; Ruzicka et al., 2020).

In this work, we used the DGRP to describe the genetic architecture of the thermal tolerance landscape in *D. melanogaster* in different thermal environments and in both sexes to understand the evolutionary potential of thermal tolerance in the face of global warming. Specifically, we used quantitative genetic analyses to estimate genetic variation, heritability, GxS, and GxE of the thermal tolerance landscape in the DGRP. Later, we identified variants and candidate genes associated with both parameters of the thermal tolerance landscape, and finally, we performed functional validation on some of the candidate genes.

## MATERIALS AND METHODS

### Drosophila stocks

One hundred isogenic lines from the Drosophila Genetic Reference Panel (DGRP) were used to perform the thermal assays. The DGRP lines were generated through 20 generations of inbreeding, which were founded using females collected from Raleigh, North Carolina, United States (MacKay et al., 2012). In our laboratory, these lines were maintained at 25°C with a photoperiod of 12L:12D and with a food medium composed of 100 g/L of fresh yeast, 80/L g of glucose, 50 g/L of wheat flour, 11 g/L of agar, 12 mL/L of nipagin, 6 mL/L of propionic acid. Newly emerged flies were separated by sex into new vials every two days. The flies used in the thermal assay were between 4 and 6 days old.

### Heat tolerance assays

For each DGRP line, five males and five females were assayed to measure their heat knockdown time at four different static temperatures: 37°C, 38°C, 39°C, and 40°C (± 0.2°C). Due to the large sampling size (100 DGRP lines x 2 sexes x 4 static temperature x 5 flies = 4,000 flies), this experiment was carried out in 12 runs. At each temperature, flies were individually placed in a well of a modified 96-well PCR plate. This plate was placed inside a climatic chamber connected to a PELT-5 thermal controller (Sable Systems International). Each trial was recorded using a web camera (Logitech), and videos were analyzed using a Python script to measure each fly’s movement (Pérez-Gálvez et al. 2023). Specifically, this script measures the changes in pixel density in each well and determines the frame where the last change in density occurs (i.e., the frame where the fly stops moving). This methodology allowed a rapid determination of the collapse time of a fly exposed to thermal stress.

### Quantitative genetic analyses of knockdown time

For each static temperature, we fitted a linear mixed model on the knockdown time to test the effect of the experimental variables on the overall variance of the model using the *lme4* package for R (Bates et al., 2015). This model included sex and experimental runs as fixed effects, and the DGRP lines and the interaction between DGRP lines and sex as random effects. Fixed effects were tested through a type-III ANOVA, whereas random effects were tested using a likelihood-ratio test implemented in the *lmerTest* package for R (Kuznetsova et al., 2017). Additionally, each model made it possible to estimate the among-line (σ^2^_L_), the line×sex (σ^2^_LS_), and the within-line (σ^2^_e_) variance components. Using these variance components, we estimated cross-sex genetic correlation separately for each temperature as *r_GMF_* = σ^2^_L_/(σ^2^_L_+ σ^2^_LS_). Following Huang et al. (2020), we also estimated the amount of line×sex variance component that is due to variation among the DGRP lines in the sign and magnitude of the difference in thermal tolerance between females and males as σ^2^_LS_ = σ_LF_ σ^2^_LM_ (1 – *r_GM_*) + (σ_LF_ – σ_LM_)^2^/2. The first term represents the contribution of changes in the rank order of lines between sexes, and the second term represents the contribution of the sex difference in the magnitude of among-line variance.

The use of DGRP lines facilitated the estimation of the broad-sense heritability of the knockdown temperature in each static temperature as *H^2^ =* σ*^2^_G_ /*(σ*^2^_G_* + σ*^2^_E_*), where σ*^2^_G_* and σ*^2^_E_* are the genetic and the environmental variances, respectively. To estimate each variance component, we used a restricted maximum likelihood approach implemented in the MTDFREML software (Boldman et al., 1993). First, we iterated the full model to estimate both variance components: the maximum likelihood value and the broad-sense heritability and its standard error. Then, to evaluate the contribution of the genetic component to the model, we compared the maximum likelihood estimated from the full model to a new maximum likelihood value calculated from a model where G was constrained to zero. This comparison was performed using a likelihood-ratio test (LRT), where the critical chi-square value was equal to 3.84 (df = 1). Genetic correlations among knockdown times assayed at different static temperatures were estimated using the same approach. The contribution of the genetic covariance to the phenotypic covariance was tested by constraining the genetic covariance to zero.

### TDT curves

We calculated the average collapse time of each static temperature, DGRP line, and sex combination. These values were regressed against the assayed static temperatures following Equation 1

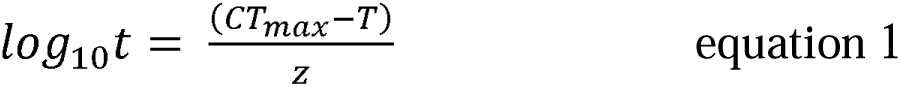

where *T* is the static temperature (°C), *CT_max_* is the maximum critical temperature (°C), t is the collapse time (min), and *z* is the thermal sensitivity (Rezende et al. 2014). These curves allowed the estimation of *CT_max_* by extrapolating the theoretical temperature where a collapse time of 1 min is obtained (log_10_ t = 0), and the estimation of *z* from the slope of each TDT line (z = -1/slope) that describes de decay in temperature tolerance following a 10-fold increase in exposure time.

We fitted linear mixed models on *CT_max_*and *z*, considering sex as a fixed effect and the DGRP lines as a random effect. The interaction between DGRP lines and sex was not included in the model because our experimental design allowed the estimation of a single value of *CT_max_* and *z* per sex and each DGRP line. We estimated the genetic and environmental variances and the broad-sense heritability of CTmax and z from the results of the linear mixed models because using the restricted maximum likelihood approach was not possible given that we only had one estimation of *CT_max_* and *z* per genotype. We estimated a CI 95% for the broad-sense heritability following Roff (1997, equation 2.28) for the standard error calculation.

### Genome-wide association analysis (GWAS)

We performed six independent GWAS for knockdown time at 37 °C, knockdown time at 38, knockdown time at 39°C, knockdown time at 40 °C, *CT_max_*, and *z*. GWAS was performed on the DGRP2 platform (http://dgrp2.gnets.ncsu.edu/) (MacKay et al., 2012). In summary, this analysis associates the phenotypic variation with biallelic variants present in the DGRP lines. The DGRP2 platform initially applies standard filtering to biallelic variants, selecting those with a minor allele frequency (MAF) greater than or equal to 0.05. Then, the phenotypic values are corrected by two criteria: (a) the *Wolbachia* infection status and (b) the effect of chromosomal inversion regions since homologous recombination does not occur in these regions (Huang et al., 2014). Finally, the corrected phenotypic values were analyzed using ANOVAs as follows: Y = µ + M +L+ E, where *Y* is the corrected value of the phenotype, M is the fixed effect of the marker (trait value difference between major and minor allele), L is the random effect of the DGRP lines, and is the residual error. Significantly associated variants (P value < 10e-5) were annotated using the FlyBase version FB5.57 (http://dgrp2.gnets.ncsu.edu/data.html). Manhattan plots were made using the R qqman package (Turner, 2014).

### Gene ontology analysis

Gene ontology (GO) enrichment analysis was performed for candidate genes associated with *CT_max_* and z. Prior to this analysis, we removed genes that do not contain any GO described in FlyBase (https://flybase.org); therefore, they are related to any functional category. The list of the filtered genes was uploaded to PANTHER version 16.0 (http://pantherdb.org), and an overrepresentation analysis was performed (PANTHER Overrepresentation Test). This analysis performs a Fisher’s exact test comparing the expected frequencies in the reference genome versus the observed frequencies of each GO category. Statistical differences between expected and observed frequencies were considered only after a False Discovery Rate correction for multiple comparisons.

### Validation of candidate genes

To functionally validate the results of GWAS, we evaluated the effect of four candidate genes (*KCNQ*, *mam*, *robo3,* and *shot*) using RNAi lines. The selection of these genes was based on the results of the gene ontology analysis, covering genes associated with the four significant categories (see Results). These RNAi lines were obtained from the Vienna Drosophila Resource Center and Bloomington Stock Center (Table S13). We used the Tub-GAL4 driver to obtain a whole-body knockdown of each gene, with the corresponding control line for each RNAi line with the same genetic background. We measured thermal tolerance on each RNAi and control line following the same procedure used for the DGRP lines, using five males and five females per genotype,temperature and replicating the whole experiment three times. The knockdown time of the experiments was analyzed in each sex and genotype by fitting linear mixed models using the *lmer* function from the lme4 R package (Bates et al., 2015) with the following model for each temperature: Y = µ + S + G + (S * G)+ R + E, where *Y* corresponds to the collapse time, *µ* corresponds to the overall mean, *S* corresponds to the fixed effect of sex, *G* corresponds to the fixed effect of the genotype, S * G corresponds to the interaction between sex and genotype, *R* corresponds to the random effect of the experimental replicate, and corresponds to the residual error.

## RESULTS

### Quantitative genetics of knockdown time

Heat knockdown time differed significantly among assayed temperatures, decreasing as the assayed temperature increased (*P value* < 2×10-16; Table S1). The average (± SD) knockdown time was 94.7 ± 26.3 min at 37°C, 52.1 ± 16.0 min at 38°C, 30.7 ± 7.8 min at 39°C, and 21.6 ± 4.5 min at 40°C (Fig. 1, 2; Table 1). The heat knockdown time was highly dependent on the genotype, sex, and assayed temperature, according to the significant interaction between these main effects in our experimental design (*P value* < 2×10-16; Table S1). Knockdown time measured at the different assayed temperatures significantly exhibited variation among DGRP lines (Table S1). The broad-sense heritability for knockdown time showed similar values across tested temperatures (range *H*^2^: 0.45 – 0.61). This result was independent of whether they were estimated using the mixed-linear model (Table 1; Table S1) or the REML approach (Table 3, Fig. S1). Genetic and environmental variances decreased with tested temperatures, but CV_L_ and CVe were relatively similar across tested temperatures (Table 1).

**Figure 1.**
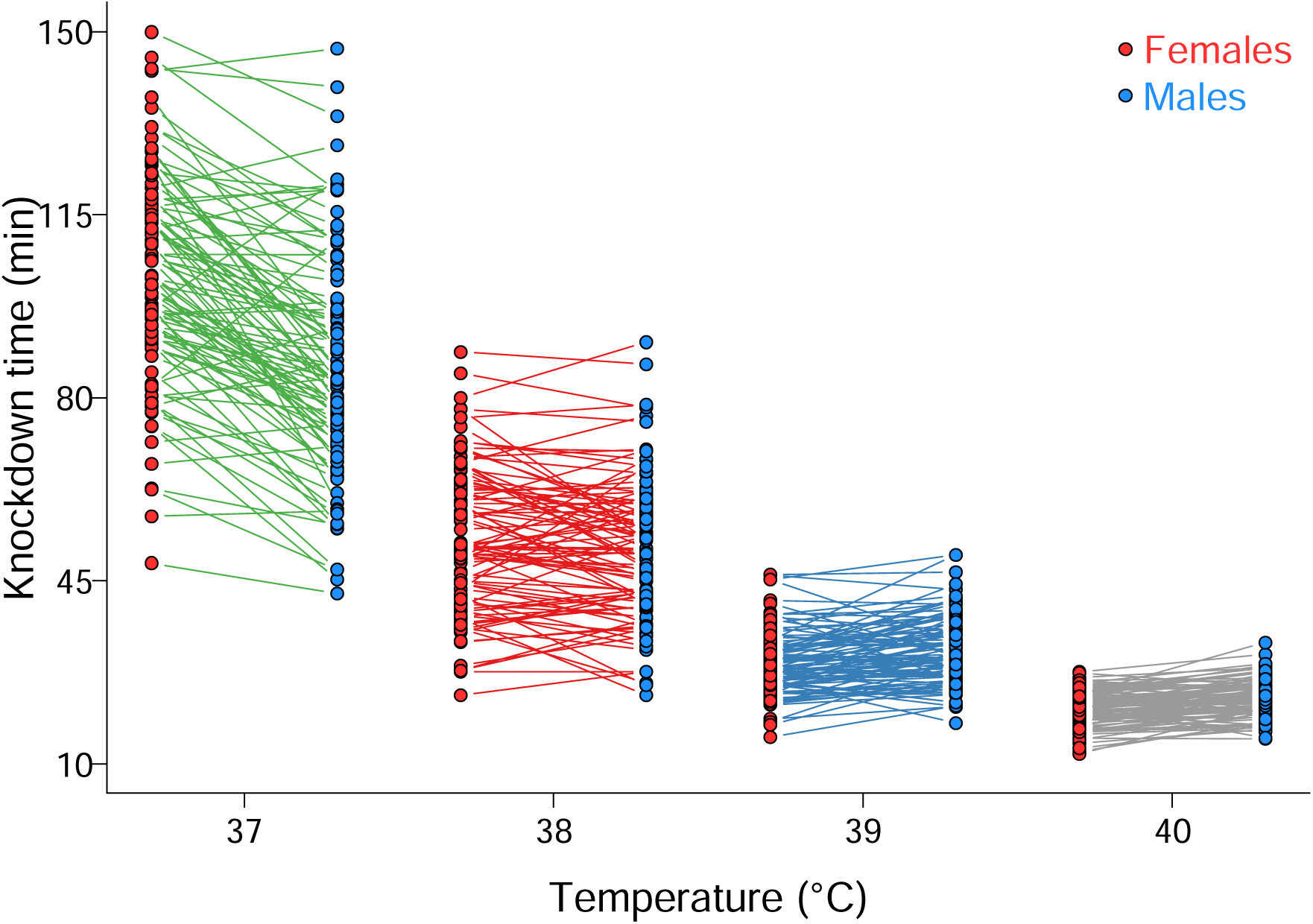
Genotype-by-sex reaction norm of the knockdown time measured in 100 DGRP lines at the different experimental temperatures. Each line corresponds to the reaction norm of a DGRP line between females (red circles) and males (blue circles).

**Table 1.** Quantitative genetic and genotype-by-sex interaction parameters for knockdown time measured at four static temperatures in the *Drosophila* Genetic Reference Panel (DGRP).

| Parameter | Static temperature |  |  |  |
| --- | --- | --- | --- | --- |
|  | 37°C | 38°C | 39°C | 40°C |
| $\bar{X}$ (females) | 103.15 min | 53.16 min | 29.36 min | 20.59 min |
| $\bar{X}$ (males) | 86.23 min | 15.57 min | 21.94 min | 22.61 min |
| $\sigma^2_L$ | 268.38 | 134.56 | 22.20 | 7.45 |
| $\sigma^2_{SL}$ | 82.97 | 23.90 | 8.87 | 2.31 |
| $\sigma^2_e$ | 243.97 | 85.64 | 24.73 | 8.13 |
| $H^2$ | 0.59 | 0.65 | 0.56 | 0.55 |
| $CV_G$ | 26.92 | 31.67 | 25.09 | 19.67 |
| $CV_e$ | 16.49 | 17.78 | 16.22 | 13.20 |
| $r_{GMF}$ | 0.764 | 0.849 | 0.715 | 0.764 |
| $\sigma_{LF}\sigma_{LM}(1 - r_{GMF})$ | 82.79 | 24.11 | 8.95 | 2.32 |
| $(\sigma_{LF} - \sigma_{LM})^2/2$ | 4.45 | 0.01 | 0.14 | 0.02 |
| % rank | 94.90 | 99.96 | 98.46 | 99.20 |

$\bar{X}$ : average value; $\sigma^2_L$ : among-line variance; $\sigma^2_{SL}$ : sex-by-line interaction variance; $\sigma^2_e$ : within-line variance; $H^2$ : broad-sense heritability calculated using the variance components estimated from the mixed linear model; $CV_G$ : coefficient of genetic variation; $CV_e$ : coefficient of environmental variation; $r_{GMF}$ : cross-sex genetic correlation; $\sigma_{LF}$ and $\sigma_{LM}$ : female and male genetic standard deviation, respectively; % rank: contribution of change in rank order of knockdown time between females and males to sex-by-line interaction variance.

**Table 2.** Genotype-by-temperature interaction parameters for knockdown time measured at four static temperatures in the *Drosophila* Genetic Reference Panel (DGRP).

| Temperature comparison | Parameter |  |  |  |
| --- | --- | --- | --- | --- |
| | $r_{GT}$ | $\sigma_{Li}\sigma_{Lj}(1 - r_{GT})$ | $(\sigma_{Li} - \sigma_{Lj})^2/2$ | % rank |
| *Females* |  |  |  |  |
| 37°C,38°C | 0.586 | 65.34 | 22.73 | 74.73 |
| 37°C,39°C | 0.370 | 58.20 | 71.45 | 44.89 |
| 37°C,40°C | 0.189 | 42.65 | 101.65 | 29.55 |
| 38°C,39°C | 0.523 | 32.06 | 26.09 | 55.14 |
| 38°C,40°C | 0.337 | 25.32 | 45.39 | 35.81 |
| 39°C,40°C | 0.549 | 7.32 | 2.66 | 73.37 |
| *Males* |  |  |  |  |
| 37°C,38°C | 0.576 | 109.34 | 28.22 | 79.22 |
| 37°C,39°C | 0.358 | 76.53 | 103.79 | 42.44 |
| 37°C,40°C | 0.213 | 51.63 | 145.33 | 26.21 |
| 38°C,39°C | 0.514 | 36.26 | 23.35 | 60.83 |
| 38°C,40°C | 0.341 | 27.09 | 44.88 | 37.64 |
| 39°C,40°C | 0.516 | 9.19 | 3.49 | 72.49 |

$r_{GT}$ : cross-temperature genetic correlation; $\sigma_{Li}$ and $\sigma_{Lj}$ : genetic standard deviation in temperatures $i$ and $j$ ; % rank: contribution of change in rank order of knockdown time between temperatures to temperature-by-line interaction variance.

**Table 3.**
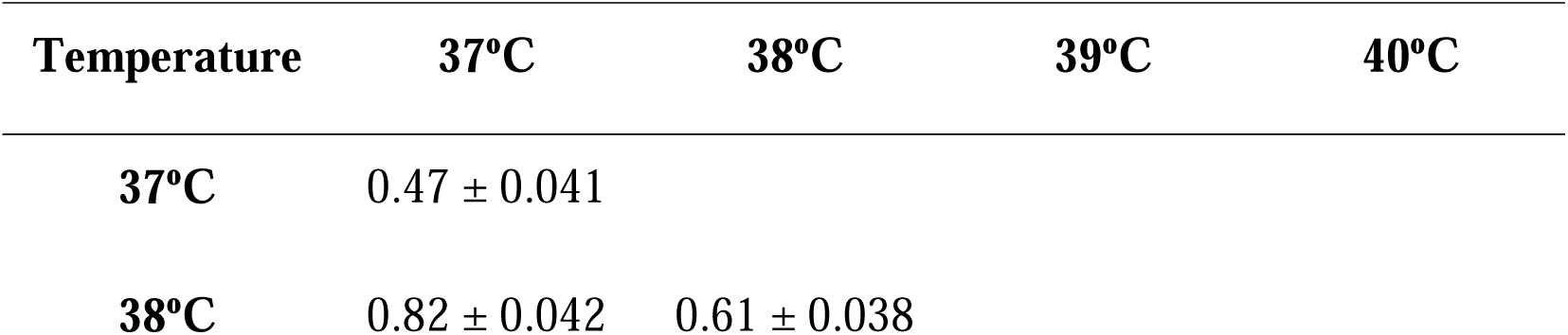
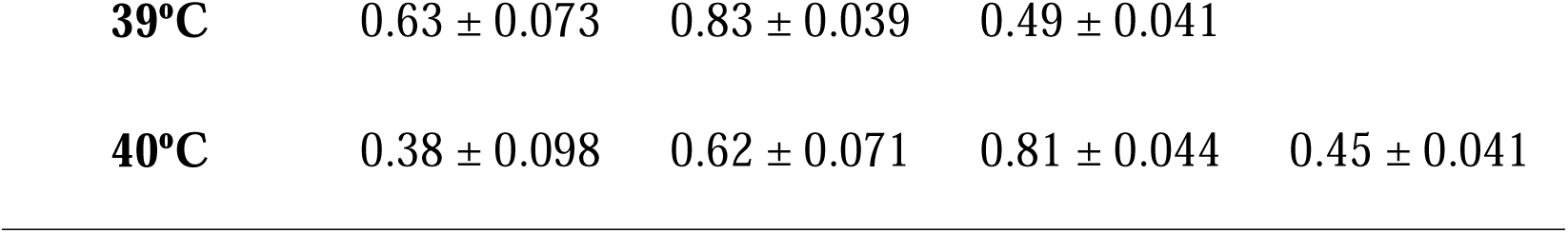
Broad-sense heritability (diagonal; ± asymptotic standard errors) and genetic correlations (off-diagonal; ± asymptotic standard errors) between heat tolerance (knockdown time) measured at different static temperatures.

| Temperature | 37°C | 38°C | 39°C | 40°C |
| --- | --- | --- | --- | --- |
| 37°C | 0.47 ± 0.041 | | | |
| 38°C | 0.82 ± 0.042 | 0.61 ± 0.038 | | |

|  |  |  |  |
| --- | --- | --- | --- |
| **39°C** | $0.63 \pm 0.073$ | $0.83 \pm 0.039$ | $0.49 \pm 0.041$ |
| **40°C** | $0.38 \pm 0.098$ | $0.62 \pm 0.071$ | $0.81 \pm 0.044$ | $0.45 \pm 0.041$ |

We found significant sex differences across tested temperatures (Table S1): females tested at 37° and 38°C showed higher tolerance than males, whereas the opposite response was observed when flies were tested at 39° and 40°C (Fig. 1, Table 1). We also found high cross-sex genetic correlations at all tested temperatures (range *r*_GME_: 0.715 – 0.849, Table 1), indicating a high correspondence for the heat tolerance in females and males of the same genotype. However, we found a significant genotype-by-sex interaction (G×S) for knockdown time at all tested temperatures (Fig. 1, Table S1). G×S is mainly explained by the changes in the rank order of the heat tolerance exhibited by females and males of the same DGRP line (range: 94.9% – 99.96%) (Table 1), which is easily visualized by the crossing of the reaction norms for knockdown time between females and males (Fig. 1).

We found significant and positive cross-temperature genetic correlations (Table 2; Table S3). Interestingly, genetic correlations decreased as the difference between tested temperatures increased. The genetic correlations were greater when estimated using the REML method than when using the variance components of the mixed linear model (Table 2, 3; Table S2). In addition, pairwise analysis between tested temperatures showed that knockdown time showed a significant genotype-by-temperature interaction (G×E), indicating that knockdown time is highly plastic across tested temperatures (Table S3, Fig. 2). However, the source of the contribution to the G×E depended on the pairs of tested temperatures compared: the closer the tested temperatures compared, the greater the contribution of the change in the rank order of heat tolerance in G×E (Table 2). This pattern was similar for both females and males.

**Figure 2.**
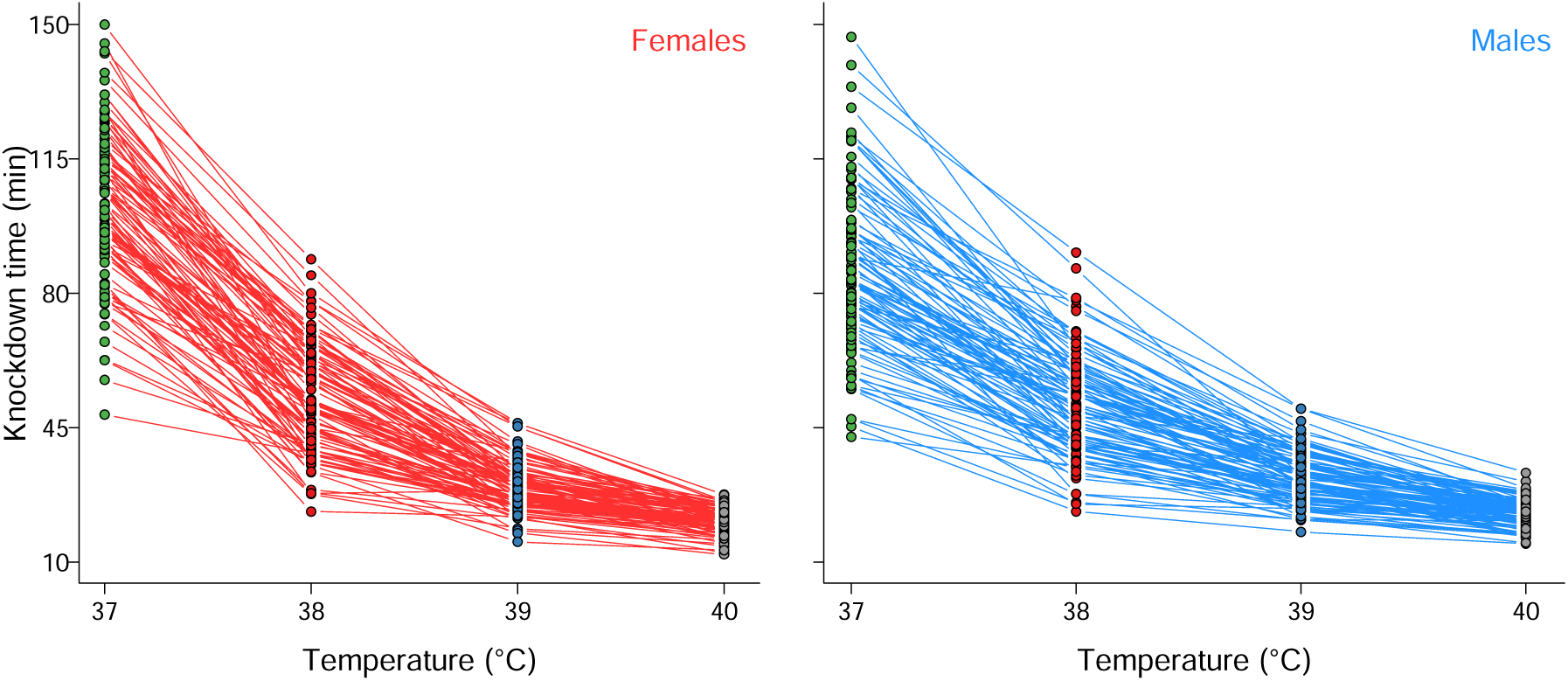
Temperature-by-sex reaction norm of the knockdown time measured in 100 of the DGRP lines for females (left) and males (right). Each line corresponds to the reaction norm of a DGRP line between two static temperatures.

### Quantitative genetics of TDT curves

We estimated 200 TDT curves (Fig. 3) with high coefficients of determination (mean *R*^2^ = 0.912, range: 0.761 – 1.000; Table S4). *CT_max_* was significantly lower for females than for males (F_1,99_ = 142.3, *P value* < 2.2×10^-2^; Fig. 3 bottom left): mean *CT_max_* for females was 45.58°C and mean *CT_max_* for males was 47.26°C. In addition, *z* values were significantly lower for females than males (F_1,99_ = 136.6, *P value* < 2.2×10^-2^; Fig. 3 bottom right): mean *z* for females was 4.38°C and mean *z* for males was 5.45°C. The TDT curves indicate that males have higher *CT_max_* than females but at the cost of exhibiting lower thermal tolerance at less extreme stress temperatures (e.g., 37°C). Furthermore, both TDT parameters showed significant genetic variation among DGRP lines (Fig. 3, Table S5). The broad-sense heritabilities for *CT_max_* and z were 0.57 (IC95% = 0.42-0.73) and 0.60 (IC95% = 0.45-0.75), respectively. On the other hand, we could not statistically test the GxS for both *CT_max_*and *z* because we only estimated one value per DGRP line and sex. However, we did observe many crossings of reaction norms for *CT_max_*and *z* between the sexes (Fig. 3).

**Figure 3.**
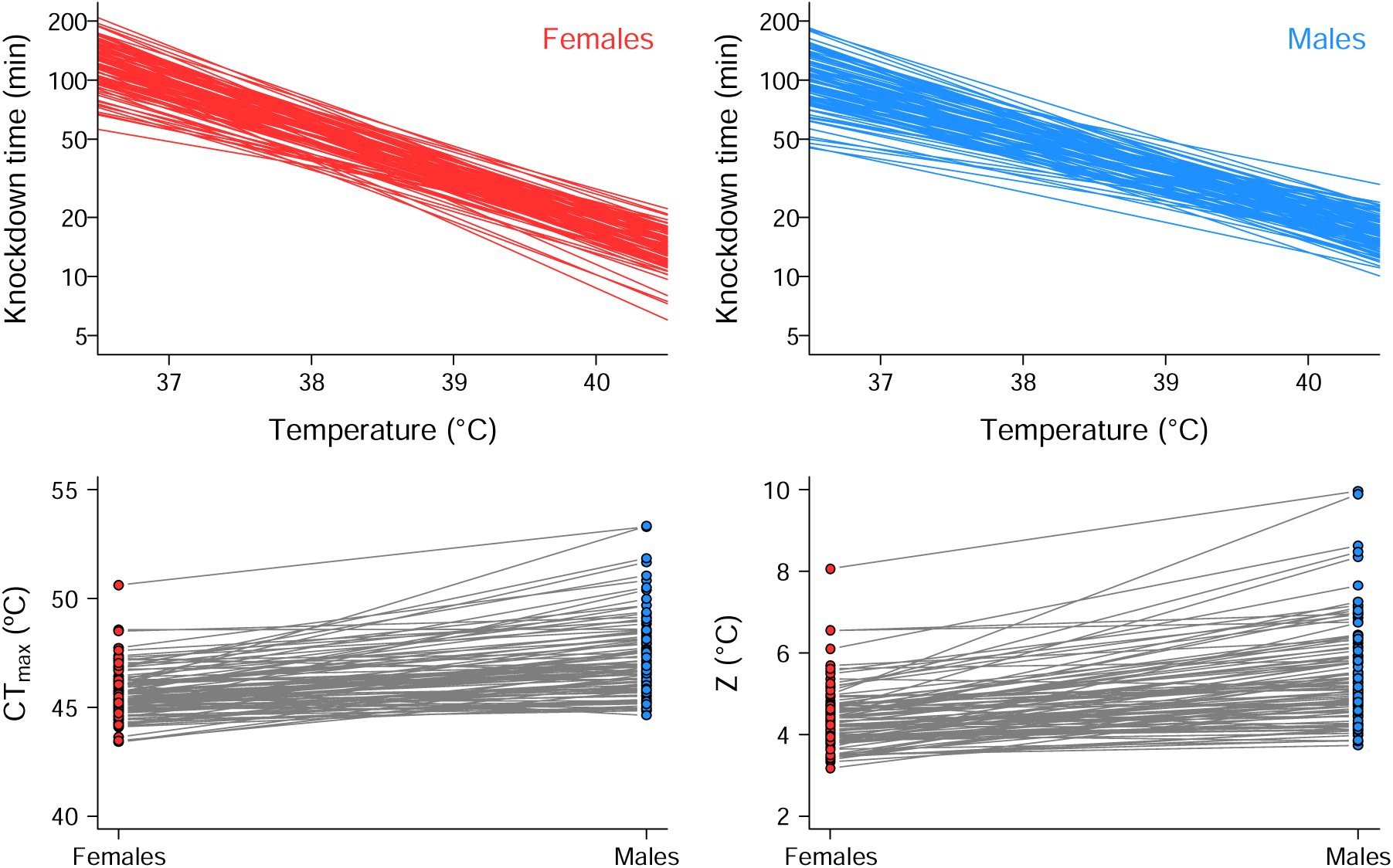
Thermal death-time curves (TDT) for females (upper left panel) and males (upper right panel) of 100 DGRP lines, where each line corresponds to the TDT curve of a DGRP line. Variations between sexes for the critical maximum temperature (CT_max_) and thermal sensitivity (z) are shown in the lower left and lower right panels, respectively.

### Gene and variants analysis

We used GWAS to identify genetic polymorphisms associated with variation in knockdown time (Table 4, Fig. S2–S5, Table S6-S9), *CT_max_*(Fig. 4; Table 5; Fig. S6), and *z* (Fig. 4; Table 5; Fig. S7). We observed that the number of variants and candidate genes associated with the average sex-pooled knockdown time increased with tested temperature (Table 4). The highest number of genetic variants associated with the knockdown time was found at the tested temperature of 40°C, mainly in chromosomes 3R and X (Fig. S5, Table S9). In addition, we found that the genetic variants and candidate genes shared between the knockdown time estimated at the different tested temperatures was very low (Table 4), suggesting that the genetic basis of heat tolerance is independent across the tested temperatures. On the other hand, the sex-pooled GWAS showed that 76 SNPs were significantly associated with *CT_max_* within or near 61 different candidate genes (Table 5, Table S10), which were distributed throughout the entire genome with a higher concentration of SNPs located on the chromosome arms 3L and 3R, and no SNPs associated with the chromosome 4 (Fig. 4). For *z*, the sex-pooled GWAS associated 140 variants that were highly concentrated in the chromosome arms 3L and 3R (Fig. 4) and mapped within or nearby 94 candidate genes (Table 5, Table S11). For these genetic variants, 51 were shared between the two TDT parameters, which mapped to 45 candidate genes (Table 5). GO analysis (Table 6; Table S12) showed an overrepresentation of biological processes related to developmental processes and cell-to-cell signaling and an overrepresentation of genes associated with the plasma membrane and cell-cell junction (cellular component category). In addition, we found that only 5 and 16 genetic variants for *CT_max_*and *z*, respectively, were shared between the sexes (Table 5).

**Figure 4.**
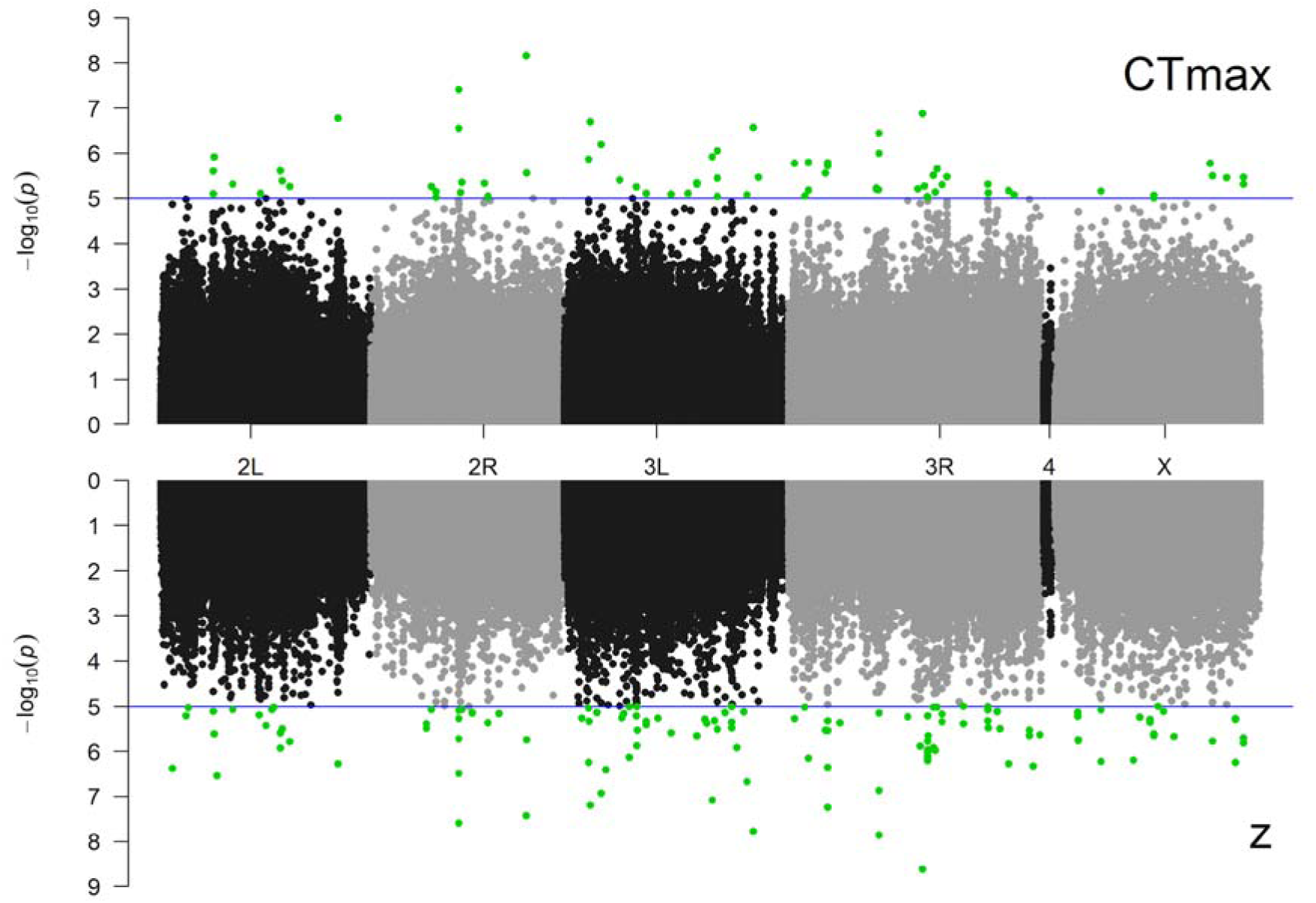
Association between genetic variants for *CT_max_* (top) and *z* (bottom) along the *Drosophila melanogaster* genome. The blue line marks the significance cutoff point at *P value* = 10^-5^, and the green circles indicate the variants with significant association with the thermal death time (TDT) curves.

**Table 4.** Results of GWAS on average sex-pooled knockdown time measured at different static temperatures in the *Drosophila* Genetic Reference Panel (DGRP). Along the diagonal, each cell indicates the number of variants associated with knockdown time and the number of candidate genes they map (parenthesis). The number of variants and candidate genes shared between knockdown time measured at different static temperatures are indicated below and above the diagonal, respectively.

| Temperature | 37°C | 38°C | 39°C | 40°C |
| --- | --- | --- | --- | --- |
| 37°C | 13 (12) | 3 | 0 | 0 |
| 38°C | 1 | 11 (13) | 0 | 1 |
| 39°C | 0 | 0 | 19 (14) | 2 |
| 40°C | 0 | 0 | 1 | 37 (29) |

**Table 5.** Results of GWAS on *CT_max_* and z (TDT parameters) in the *Drosophila* Genetic Reference Panel (DGRP). Along the diagonal, each cell indicates the number of variants associated with each trait and the number of candidate genes to which they map (parenthesis). The number of variants and candidate genes shared between knockdown time measured at different static temperatures are indicated below and above the diagonal, respectively.

| Parameter | $CT_{max}$ | $z$ | $CT_{max}$ | $CT_{max}$ | $z$ | $z$ |
| --- | --- | --- | --- | --- | --- | --- |
|  |  |  | females | males | females | males |
| $CT_{max}$ | 76 (61) | 45 | 15 | 28 | 18 | 36 |
| $z$ | 51 | 140 (94) | 26 | 22 | 40 | 49 |
| $CT_{max}$ females | 20 | 40 | 106 (68) | 8 | 51 | 8 |
| $CT_{max}$ males | 32 | 24 | 5 | 59 (45) | 8 | 31 |
| $z$ females | 20 | 64 | 81 | 6 | 232 (145) | 11 |
| $z$ males | 45 | 63 | 10 | 45 | 16 | 91 (62) |

**Table 6.**
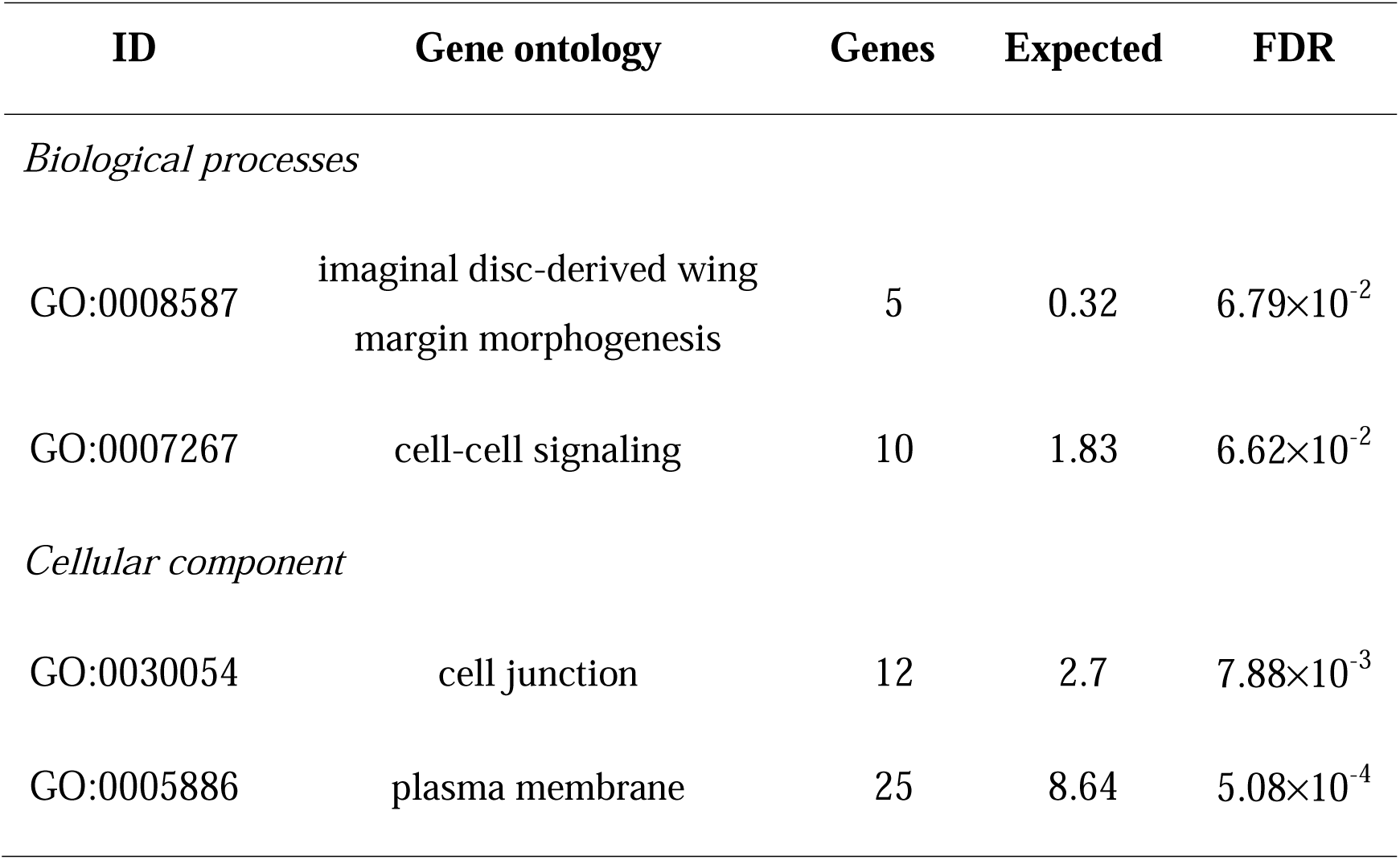
Result of gene ontology enrichment analysis of candidate genes (PANTHER Overrepresentation Test).

| ID | Gene ontology | Genes | Expected | FDR |
| --- | --- | --- | --- | --- |
| *Biological processes* |  |  |  |  |
| GO:0008587 | imaginal disc-derived wing margin morphogenesis | 5 | 0.32 | $6.79 \times 10^{-2}$ |
| GO:0007267 | cell-cell signaling | 10 | 1.83 | $6.62 \times 10^{-2}$ |
| *Cellular component* |  |  |  |  |
| GO:0030054 | cell junction | 12 | 2.7 | $7.88 \times 10^{-3}$ |
| GO:0005886 | plasma membrane | 25 | 8.64 | $5.08 \times 10^{-4}$ |

### Effect size of variants and validation of candidate genes

We observed that most of the variants associated with *CT_max_* had a negative effect size (i.e., homozygous flies for the minor allele showed higher *CT_max_* than homozygous flies for the major allele). In contrast, all variants associated with *z* had a negative effect size (i.e., homozygous flies for the minor allele showed higher z values than flies homozygous for the major allele and, therefore, are less sensitive to change of temperature) (Fig. 5-6). We also observed that the effect size of these variants was inversely proportional to the frequency of the minor allele; thus, alleles that increase the thermal tolerance are at low frequency in the DGRP lines studied (Fig. 5).

**Figure 5.**
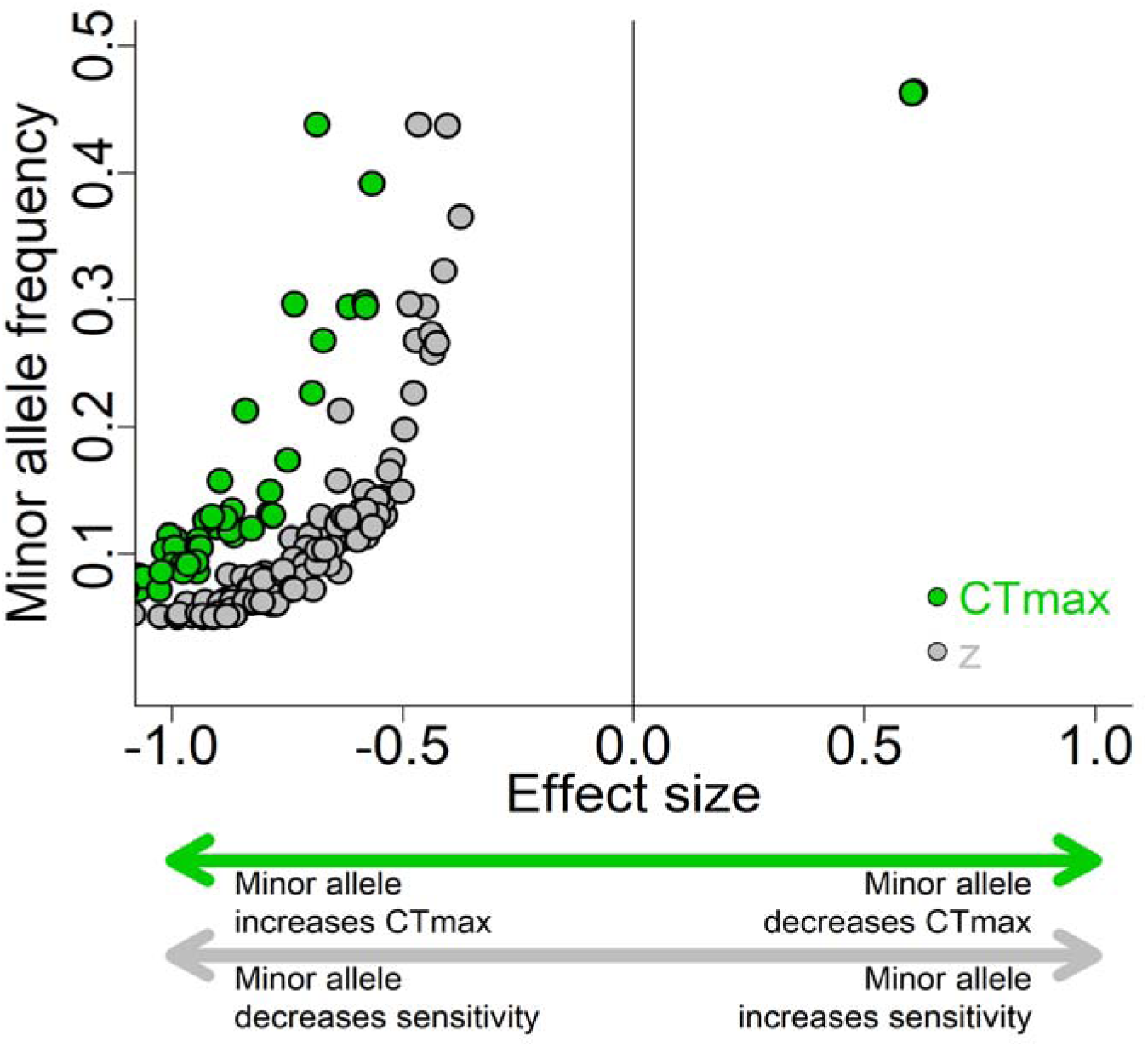
Relationship between the minor allele frequency and effect size of variants significantly associated with the thermal-death-time (TDT) parameters: the critical maximum temperature (*CT_max_*, green circles) and the thermal sensitivity (*z*, gray circles).

**Figure 6.**
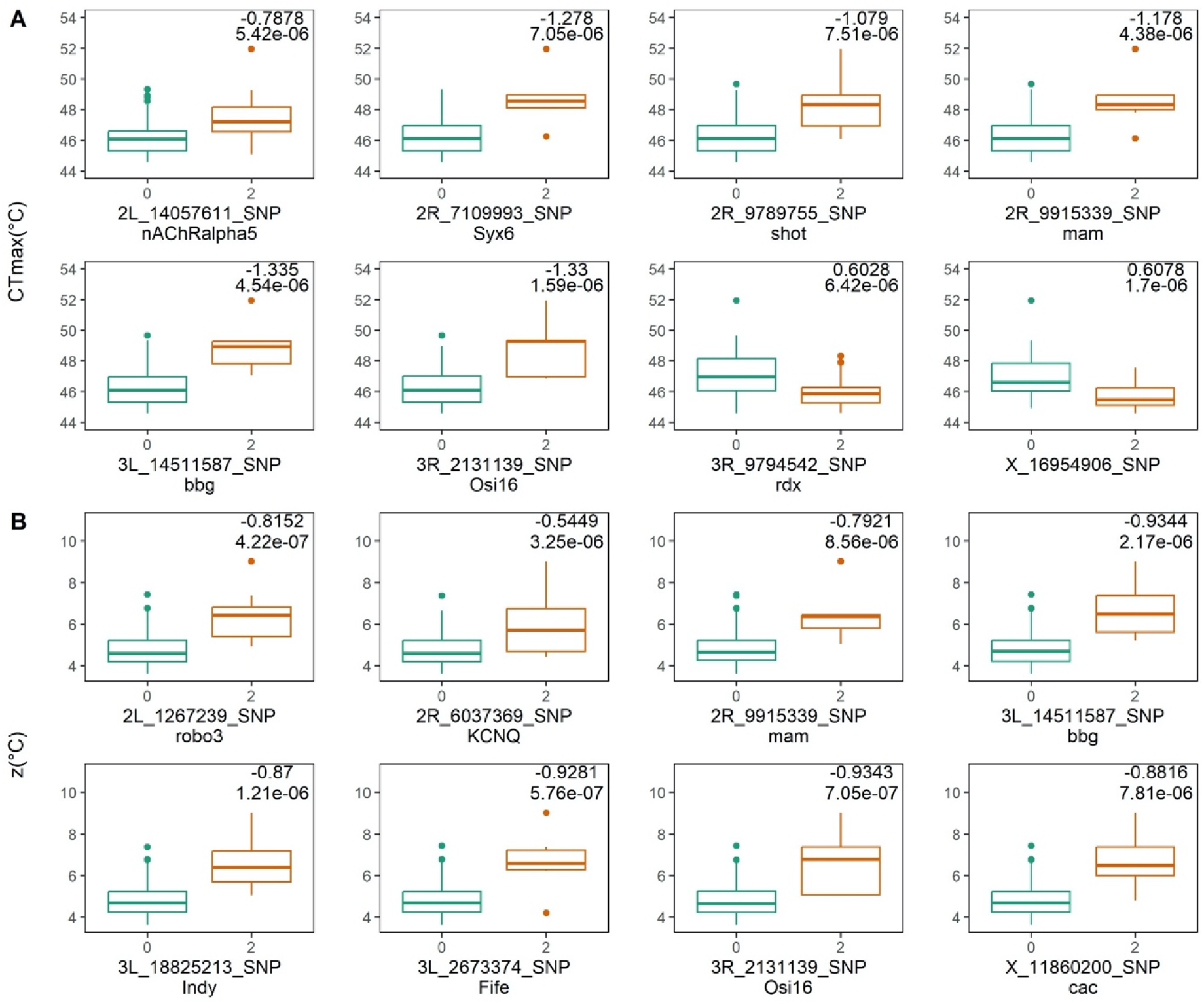
The effect of some variants is significantly associated with the *CT_max_* (A) and *z* (B). Each plot shows the phenotype distribution for the homozygous flies for the major allele (0) and the homozygous flies for the minor allele (2) for some variants with a larger effect size identified in the GWAS analyses. The X-axis shows the name of the variant and the gene it maps. Numbers in the upper right corners show the effect size (upper) and the *P value* of the comparison (lower).

Regarding gene validation, we found that the RNAi-*mam* line had a higher thermal tolerance than the control line at 38°C in both sexes and at 39°C only in males (Fig. 7, Table S14). On the other hand, the RNAi-*KCNQ* line showed a lower thermal tolerance than the control line at 38°C and 39°C only in males, while the RNAi-*robo3* line showed a lower thermal tolerance than the control lines at 40°C only in males. No difference was found between RNAi-*shot* and control lines. Finally, we found no difference between control and RNAi lines on *CT_max_*and *z* parameters (Table S14). This suggests that the candidate genes tested only affect the knockdown time at a specific tested temperature and in a sex-specific manner but do not affect the TDT curves.

**Figure 7.**
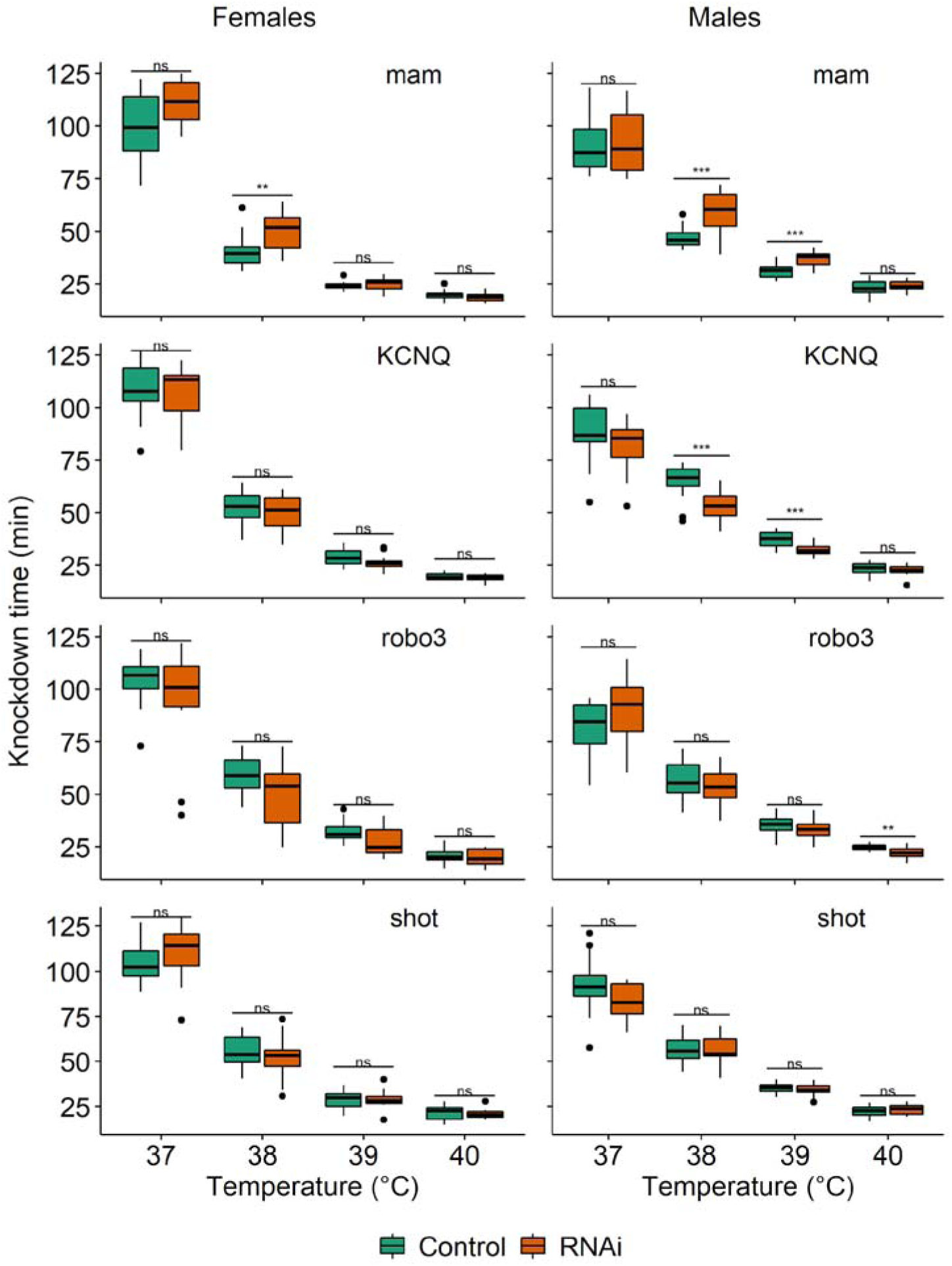
RNAi-mediated knockdown of candidate genes associated with thermal-death-time (TDT) curves for females (left) and males (right). Symbols above boxplots represent the statistical results of the comparisons of the knockdown time between control (green) and RNAi (orange) lines: ns: non-significant, *: *P value* < 0.05, **: *P value* < 0.01; ***: *P value* < 0.001.

## DISCUSSION

This work reveals the genetic basis of the thermal tolerance landscape in *D. melanogaster*, which has a genetic component that can be quantified and identified through quantitative genetic and genomic analyses. We also found that heat tolerance is influenced by G×E and G×S, suggesting that the genetic architecture of heat tolerance depends on multiple factors. Our findings indicate that heat tolerance is genetically determined; however, the specific genes related to heat tolerance vary depending on the temperature being tested. The main novelty of the present study is that we combined a resource panel with high genetic variations and the thermal tolerance landscape (TDT curves) to explore the genetic architecture of thermal tolerance to high temperatures in a classic genetic and evolutionary model such as *D. melanogaster*. Our results demonstrate that heat tolerance has enough potential to evolve under different natural selection scenarios but that its genetic variation can be maintained due to G×E and G×S mechanisms. The genetic basis of heat tolerance changes depending on the thermal environment.

### Evolutionary potential of the thermal tolerance landscape

Broad-sense heritabilities and genetic and environmental coefficients of variation for knockdown time were similar across the tested temperatures, indicating that this trait could undergo selection-driven evolution regardless of the thermal environment, and even though the true heritability (i.e., narrow sense heritability) is predicted to be lower than the currently reported values. However, similar heritability values across tested temperatures were unexpected because previous studies had stated that longer experiments tend to increase the effect of confounding experimental variables (e.g., starvation, desiccation) that increase the environmental variation, resulting in lower estimates of heritabilities than shorter assays (Mitchell & Hoffmann, 2010; Overgaard et al., 2011; Rezende et al., 2011). Here, the longest trial (37°C) was, on average, 4-5 times longer than the shortest trial (40°C), which should have affected the heritability estimates of the knockdown time. However, Rezende et al., (2011) described the effect of desiccation and starvation are important for thermal tolerance when the tests exceed 140 minutes; thus, the influence of these confounding variables should be low for all the tests carried out in the present work (the longest tests lasted an average of 95 minutes).

We also found positive genetic correlations in heat tolerance across assay temperatures, but the magnitude of the genetic correlation decreased as the difference between assay temperatures increased. This pattern has been previously described for cold tolerance in *D. melanogaster* (Ørsted et al. 2019), life-history traits in *D. serrata* (Stinchcombe et al. 2010, and body size in the nine-spined stickleback (*Pungitius pungitius*) (Fraimout et al. 2022). Delving into greater detail, Ørsted et al. (2019) suggested that cold tolerance has a shared genetic basis, but the underlying genes become more different as the thermal environments diverge. Here, we found that the genetic variants and candidate genes shared between the knockdown time estimated at the different assay temperatures was very low, suggesting that the genetic basis of heat tolerance is independent across the assay temperatures. This mechanism, known as environment-dependent gene action, can have significant consequences on the evolutionary prediction based on genetic information because selection target genes can change depending on the environmental conditions (Ørsted et al. 2019). We also found a significant genotype-environment interaction (GxE) for the knockdown time, which means that the genotypes respond differently to the thermal challenge depending on the exposure temperature. Other studies have also found significant GxE for thermal traits such as cold tolerance (Ørsted et al. 2019), cold and heat hardiness (Freda et al. 2019). This genetic variation in the phenotypic plasticity of the thermal tolerance should contribute to the adaptive responses to different thermal environments and maintain the genetic diversity in natural populations.

From the TDT curves, our estimates of *CT_max_* and its heritability were higher than those described in previous forecasts for heat tolerance in the DGRP (Lecheta et al., 2020; Rolandi et al., 2018). However, these two studies used dynamic assays and we used static assays, which are known to estimate higher heritability values than dynamic assays (Castañeda et al., 2019; Mitchell & Hoffmann, 2010). The *CT_max_* values are more in line with what was reported by Jørgensen et al. (2021), who also used the TDT curves to estimate the thermal limits. Some studies indicate that thermal tolerance has a limited evolutionary capacity (Kellermann et al., 2012; Kelly et al., 2012), while others support the opposite (Folk et al., 2006; Geerts et al., 2015; Logan & Cox, 2020). The results of this work using a large number of genotypes indicate heritability in the parameters of the thermal tolerance landscape; therefore, they support the idea that ectotherm populations could potentially evolve in the face of thermal changes in their environment. This is supported by laboratory evolutionary experiments where organisms exposed to thermal selection undergo rapid evolutionary changes in their thermal limits (Hangartner & Hoffmann, 2016; Mesas et al., 2021; Sambucetti et al., 2010).

### Genes and variants associated with the thermal tolerance landscape

Many genetic variants associated with the parameters of the thermal tolerance landscape were identified in the present study. Most of these candidate variants are located in non-coding regions such as introns and UTR regions. Genetic variation in non-coding regions has been related to the regulatory processes of gene expression and alternative splicing processes, which could affect the transcriptional plasticity and the adaptive response of populations inhabiting changing environments (Kelly, 2019; A. Li et al., 2021). Considering this, it is unsurprising that several of our candidate genes exhibit differential expression between high and low latitudes populations of *Drosophila* (Zhao et al., 2015). The differential gene expression is an important mechanism exhibited by populations exposed to stressful environments because adaptive plasticity may facilitate evolutionary rescue in these environments and “buying time” for organisms to display a phenotypic response capable of reducing the impact of changing environments on natural populations (Diamond & Martin, 2021).

According to the genetic variants, they are mapped to candidate genes related to processes occurring on the periphery of the cell, such as communication between cells. For instance, we found some genes participating in synaptic processes, such as *nAChRalpha5*, which encodes receptor subunits nicotinic acetylcholine (Lansdell et al., 2012); *fife*, which is associated with the neurotransmitter release (Bruckner et al., 2012); *syx6*, which is believed to be an integral component of synaptic vesicles (Gaudet et al., 2011); and *cac* that encodes part of the presynaptic voltage-gated calcium channels (Chang et al., 2014). These genetic associations indicate a relationship between the motor capacity of the flies and the variation observed in *CT_max_* and *z*. Additionally, we also found a subset of candidate genes that are important during the fly development, such as *mam* and *dl*, which participate in the pathway Notch (Ahimou et al., 2004; Gomez-Lamarca et al., 2018); and *sgg*, which is involved in the canonical *Wnt* signaling pathway (Miech et al., 2008). Among this category, only the *fid* gene has been related to thermal tolerance, which encodes a protein that participates in thermal resistance during the larval stage of *Drosophila* (Honjo et al., 2016). Therefore, other candidate genes involved in larval development may impact the thermal capacity of adults.

We also found that most of the variants were positively associated with the thermal tolerance landscape, increasing the maximum critical temperature, and decreasing thermal sensitivity. Additionally, we found an inversely proportional relationship between the effect size and the frequency of the minor allele. Lecheta et al. (2020) described this pattern in the genetic polymorphisms for *CT_max_*and this coincides with other phenotypes described for the DGRP (MacKay et al., 2012; Weber et al., 2012). These results suggest that these genes are under selection in natural populations since variants that negatively affect the thermal tolerance landscape would be rapidly eliminated from the population through purifying selection. On the other hand, alleles that have a large positive effect on thermal tolerance are present in low frequency, which may indicate that these alleles are under purifying selection by interacting negatively with another fitness trait (Barton & Keightley, 2002). Considering that both parameters of the thermal tolerance landscape are correlated and, therefore, share many of the variants and candidate genes, it is expected that both the maximum critical temperature and thermal sensitivity could evolve together in populations.

Interestingly, 25% of our candidate genes (27 out of 110) have been described to show latitudinal or temporal variation in previous studies on natural populations of *D. melanogaster* (Table S15). For example, four of these genes (*eip63E*, *ino80*, *MCO3*, and *egg*) showed allelic variation along a latitudinal gradient on the East Coast of the United States (Fabian et al., 2012; Machado et al., 2016). Another set of candidate genes reported in the present study showed latitudinal variation in Australia (CG7720 and *Dlc90F*) (Kolaczkowski et al., 2011) and Europe (*mam*, *MCO3*, *corn*, and *sug*) (Kapun et al., 2020). We also found that some candidate genes related to the thermal tolerance landscape showed temporal variation; for example, the genes *sgg*, *Eip74EF*, *tei*, *cac*, CG7737, and CG34354 showed positive selection between summer and autumn in a North American population (Rudman et al., 2022). Finally, several of our candidate genes showed differential expression patterns between populations from temperate and tropical climates, including genes mentioned previously, like *cac* and *fid* (Zhao et al., 2015). Therefore, we found that some candidate genes associated with thermal tolerance in the present study also show variation among natural populations. This suggests that our candidate genes are under selection due to their role in the adaptive response to different thermal environments.

It is important to mention that none of the candidate genes reported here coincide with previous ones reported in DGRP studies estimating the thermal tolerance using dynamic assays (Lecheta et al., 2020; Rolandi et al., 2018). However, both studies found genetic variants associated with developmental functions, which supports the idea that gene tuning during development could affect the capacity to withstand thermal stressors in adults.

Similarly, no association with *hsp* genes was found to explain the genetic variability in the thermal tolerance landscape of DGRP lines. The proteins of the HSP family have a highly conserved sequence and structure in nature (Desai et al., 2010), and the DGRP population likely has low genetic variability in the genes encoding these proteins. Although these genes are important in the physiological response to thermal challenges, genetic variation could not be at the SNP level. Indeed, *hsp70* gene expression is positively correlated with *hsp70* gene copy number, which confers a higher thermotolerance in *D. melanogaster* (Bettencourt et al., 2008).

### Sexual dimorphism of the thermal tolerance landscape

Several works have studied the sex difference in the thermal tolerance of *Drosophila*. Some authors found no significant differences between the sexes (Van Heerwaarden et al., 2016), others found higher tolerance for females (Mitchell & Hoffmann, 2010), while others reported higher tolerance for males (Lecheta et al., 2020; Castañeda et al., 2015). However, something in common among these studies is they used a single static temperature to estimate the thermal tolerance. Here, we observed that the thermal tolerance difference between sexes depends on the temperature used in the assay. If static tests are carried out at mildly stressful temperatures (<37°C), females exhibit a higher tolerance than males.

Conversely, if extreme temperatures (>40°C) are used, males have a higher tolerance than females. At some temperatures, no differences between sexes were found, as in the case of 38°C. This is interesting because this temperature is one of the temperatures most frequently used in the static assays (Blackburn et al., 2014; Castañeda et al., 2019; Mitchell & Hoffmann, 2010; Van Heerwaarden et al., 2016). Therefore, it is important to consider the temperature used in the static assays if the study is seeking to evaluate the sexual dimorphism of thermal tolerance. Additionally, the genetic basis of the thermal tolerance landscape appears to be sex-specific, given that few variants and candidate genes were shared between sexes for the parameters of *CT_max_*and *z*.

Future studies could focus exclusively on this sexual dimorphism of the thermal tolerance landscape and even study the existence of genetic determinants that could have an antagonistic pleiotropic effect between sexes. The differences in the thermal tolerance landscape between the sexes could have important repercussions in natural populations.

Climate change generates thermal changes associated with two different selective pressures: on the one hand, it generates an increase in the global average temperature, and on the other hand, it generates an increase in thermal variability through more frequent extreme events (Angilletta, 2009). Considering this context, females would have a better adaptive capacity to the first component, given their greater capacity to resist mild stressful temperatures. Meanwhile, males could better adapt to more variable thermal conditions, given their lower thermal sensitivity and better resistance to extreme temperatures. Therefore, the sexual dimorphism observed in the thermal tolerance landscape may arise from the distinct ways thermal selective pressures affect males and females.

## CONCLUSIONS

Studying populations’ ability to cope with environmental thermal stress through phenotypic plasticity or adaptive evolution is crucial for understanding present and future shifts in species distribution and survival. In this context, the DGRP is a robust tool to study the genetic underpinnings of the thermal tolerance landscape. Our results suggest that the genetic variation and phenotypic plasticity of the thermal tolerance should contribute to the adaptive response of *D. melanogaster* to environmental change. Additionally, the thermal tolerance landscape is characterized by a sexual dimorphism, which may confer advantages to females in a particular thermal environment. In contrast, males can exhibit an advantage in a different environment. Future efforts are needed to study how these findings impact natural populations; however, here, we can suggest that some genes showing spatial and/or temporal variation contribute to the thermal tolerance in *Drosophila*.

## ACKNOWLEDGMENTS

We thank Noemi Candia and Gonzalo Olivares for their assistance in fly manipulation and stock maintenance. We also thank Nicholas M. Teets, David N. Awde, and Fernan Pérez-Gálvez for their advice on using the script for the video analysis.

## AUTHOR CONTRIBUTIONS

JS and LEC conceptualized the study. JS performed the experiments, analyzed the videos, and performed statistical and genomic analysis. LEC performed a quantitative genetic analysis. JS wrote the first draft of the manuscript, which LEC reviewed and edited. PO facilitated the flies for the study and supervised the functional validation of candidate genes. FP contributed to the literature review and discussion of the study.

## DATA AVAILABILITY STATEMENT

Data are available on Figshare and the link is provided for the review process. Codes for analyses and figures are available on Github (https://github.com/JSlox/TTL-DGRP).

## Supplementary information

Table S1. Results of the linear mixed models performed on the knockdown time data for each tested temperature.

Table S2. Genetic variances (diagonal) of the heat tolerance (knockdown time) measured at 37, 38, 38 and 40°C, and their respective genetic covariances (off-diagonal). The statistical contribution of the genetic component was tested using likelihood-ratio tests with one degree of freedom (chi-square values are presented in parenthesis with: \**P value* < 0.05, ** *P value* < 0.01, and *** *P value* < 0.001).

Table S3. Results of the linear mixed models performed on the knockdown time data between temperatures.

Table S4. Knockdown time per temperature, CTmax and z estimates, and coefficient of correlation (r2) between *CT_max_* and thermal sensitivity (*z*) for the 100 DGRP lines studied, separated by sex.

Table S5. Results of the linear models performed on *CT_max_* and thermal sensitivity (*z*).

Table S6. Candidate variants associated with knockdown time at 37°C, genes to which they map, and the location of the variant. Results for the average (A) sex-pooled analysis, female (F) analysis and male (M) analysis.

Table S7. Candidate variants associated with knockdown time at 38°C, genes to which they map, and the location of the variant. Results for the average (A) sex-pooled analysis, female (F) analysis and male (M) analysis.

Table S8. Candidate variants associated with knockdown time at 39°C, genes to which they map, and the location of the variant. Results for the average (A) sex-pooled analysis, female (F) analysis and male (M) analysis.

Table S9. Candidate variants associated with knockdown time at 40°C, genes to which they map, and the location of the variant. Results for the average (A) sex-pooled analysis, female (F) analysis and male (M) analysis.

Table S10. Candidate variants associated with *CT_max_*, genes to which they map and the location of the variant.

Table S11. Candidate variants associated with thermal sensitivity (*z*), genes to which they map, and the location of the variant.

Table S12. Genes corresponding to each gene ontology category significantly overrepresented in the candidate genes for *CT_max_* and thermal sensitivity (*z*).

Table S13. *Drosophila* stocks used in functional validation.

Table S14. Results of ANOVA performed on the knockdown time data of RNAi flies. Significant codes: \*\*\**P values* < 0.00025, ** *P values* < 0.0025 and * *P values* < 0.0125.

Table S15. Candidate genes that show latitudinal or temporal variation in previous works in *Drosophila melanogaster*.

## REFERENCES

Ahimou, F., Mok, L., Bardot, B., & Wesley, C. (2004). The adhesion force of Notch with Delta and the rate of Notch signaling. Journal of Cell Biology, 167(6), 1217–1229. 10.1083/JCB.200407100

Anderson, A., Collinge, J., Hoffmann, A., Kellett, M., & McKechnie, S. (2003). Thermal tolerance trade-offs associated with the right arm of chromosome 3 and marked by the hsr-omega gene in Drosophila melanogaster. Heredity, 90(2), 195–202. 10.1038/sj.hdy.6800220

Angilletta, M. J. (2009). Temperature and the Life History. En M. J. Angilletta Jr. (Ed.), Thermal Adaptation: A Theoretical and Empirical Synthesis (p. 0). Oxford University Press. 10.1093/acprof:oso/9780198570875.003.0006

Araújo, A., Reis, M., Rocha, H., Aguiar, B., Morales-Hojas, R., Macedo-Ribeiro, S., Fonseca, N., Reboiro-Jato, D., Reboiro-Jato, M., Fdez-Riverola, F., Vieira, C., & Vieira, J. (2013). The Drosophila melanogaster methuselah Gene: A Novel Gene with Ancient Functions. PLoS ONE, 8(5), 63747. 10.1371/journal.pone.0063747

Barton, N., & Keightley, P. (2002). Understanding quantitative genetic variation. Nature Reviews Genetics 2001 3:1, 3(1), 11-21. 10.1038/nrg700

Bates, D., Mächler, M., Bolker, B., & Walker, S. (2015). Fitting linear mixed-effects models using lme4. Journal of Statistical Software, 67(1), 1–48. 10.18637/jss.v067.i01

Beitinger, T., & Lutterschmidt, W. (2011). Temperature | Measures of Thermal Tolerance. En Encyclopedia of Fish Physiology (Vol. 3, pp. 1695–1702). Elsevier Inc. 10.1016/B978-0-12-374553-8.00200-8

Bettencourt, B. R., Hogan, C. C., Nimali, M., & Drohan, B. W. (2008). Inducible and constitutive heat shock gene expression responds to modification of Hsp70 copy number in Drosophila melanogaster but does not compensate for loss of thermotolerance in Hsp70null flies. BMC Biology, 6(1), 5. 10.1186/1741-7007-6-5

Blackburn, S., Van Heerwaarden, B., Kellermann, V., & Sgro, C. (2014). Evolutionary capacity of upper thermal limits: Beyond single trait assessments. Journal of Experimental Biology, 217(11), 1918–1924. 10.1242/JEB.099184/257307/AM/EVOLUTIONARY-CAPACITY-OF-UPPER-THERMAL-LIMITS

Boldman, K., Kriese, L. A., Van Vleck, L., Tassell, C. P., & Kachman, S. D. (1993). A Manual for use of MTDFREML: A set of programs to obtain estimates of variances and covariances. [Draft]. USDA.

Bruckner, J., Gratz, S., Slind, J., Geske, R., Cummings, A., Galindo, S., Donohue, L., & O’Connor-Giles, K. (2012). Fife, a Drosophila Piccolo-RIM Homolog, Promotes Active Zone Organization and Neurotransmitter Release. Journal of Neuroscience, 32(48), 17048–17058. 10.1523/JNEUROSCI.3267-12.2012

Castañeda, L., Rezende, E., & Santos, M. (2015). Heat tolerance in Drosophila subobscura along a latitudinal gradient: Contrasting patterns between plastic and genetic responses. Evolution, 69(10), 2721–2734. 10.1111/EVO.12757

Castañeda, L., Romero Soriano, V., Mesas, A., Roff, D., & Santos, M. (2019). Evolutionary potential of thermal preference and heat tolerance in Drosophila subobscura. Journal of Evolutionary Biology, 32(8), 818–824. 10.1111/jeb.13483

Chang, J., Hazelett, D., Stewart, J., & Morton, D. (2014). Motor neuron expression of the voltage-gated calcium channel cacophony restores locomotion defects in a Drosophila, TDP-43 loss of function model of ALS. Brain Research, 1584, 39-51. 10.1016/J.BRAINRES.2013.11.019

Chown, S., Jumbam, K., Sørensen, J., & Terblanche, J. (2009). Phenotypic variance, plasticity and heritability estimates of critical thermal limits depend on methodological context. Functional Ecology, 23(1), 133–140. 10.1111/j.1365-2435.2008.01481.x

Connallon, T., & Clark, A. (2013). Evolutionary inevitability of sexual antagonism. Proceedings of the Royal Society B: Biological Sciences, 281(1776). 10.1098/rspb.2013.2123

Dahlgaard, J., Loeschcke, V., Michalak, P., & Justesen, J. (1998). Induced thermotolerance and associated expression of the heat-shock protein Hsp70 in adult *Drosophila melanogaster*. Functional Ecology, 12(5), 786–793. 10.1046/j.1365-2435.1998.00246.x

Delclos, P. J., Adhikari, K., Hassan, O., Cambric, J. E., Matuk, A. G., Presley, R. I., Tran, J., Sriskantharajah, V., & Meisel, R. P. (2021). Thermal tolerance and preference are both consistent with the clinal distribution of house fly proto-Y chromosomes. Evolution Letters, 5(5), 495–506. 10.1002/EVL3.248

Desai, N., Agarwal, A., & Uplap, S. (2010). HSP: Evolved and conserved proteins, structure and sequence studies. International Journal of Bioinformatics Research, 2(2), 67–87. 10.9735/0975-3087.2.2.67-87

Deutsch, C., Tewksbury, J., Huey, R., Sheldon, K., Ghalambor, C., Haak, D., & Martin, P. (2008). Impacts of climate warming on terrestrial ectotherms across latitude. Proceedings of the National Academy of Sciences of the United States of America, 105(18), 6668–6672. 10.1073/pnas.0709472105

Diamond, S. E., & Martin, R. A. (2021). Buying Time: Plasticity and Population Persistence. En Phenotypic Plasticity & Evolution. CRC Press.

Fabian, D. K., Kapun, M., Nolte, V., Kofler, R., Schmidt, P. S., Schlötterer, C., & Flatt, T. (2012). Genome-wide patterns of latitudinal differentiation among populations of Drosophila melanogaster from North America. Molecular Ecology, 21(19), 4748–4769. 10.1111/J.1365-294X.2012.05731.X

Fedra, P. J., Ali, Z. M., Heter, N., Ragland, G. J., & Morgan, T. J. (2019). Stage-specific genotype-by-environment interaction for cold and heat hardiness in *Drosophila melanogaster*. Heredity, 123: 479–491. 10.1038/s41437-019-0236-9

Folk, D., Zwollo, P., Rand, D., & Gilchrist, G. (2006). Selection on knockdown performance in Drosophila melanogasterimpacts thermotolerance and heat-shock response differently in females and males. Journal of Experimental Biology, 209(20), 3964–3973. 10.1242/JEB.02463

Gaudet, P., Livstone, M., Lewis, S., & Thomas, P. (2011). Phylogenetic-based propagation of functional annotations within the Gene Ontology consortium. Briefings in Bioinformatics, 12(5), 449–462. 10.1093/BIB/BBR042

Geerts, A., Vanoverbeke, J., Vanschoenwinkel, B., Van Doorslaer, W., Feuchtmayr, H., Atkinson, D., Moss, B., Davidson, T., Sayer, C., & De Meester, L. (2015). Rapid evolution of thermal tolerance in the water flea Daphnia. Nature Climate Change 2015 5:7, 5(7), 665-668. 10.1038/nclimate2628

Giannakou, M., Goss, M., Jünger, M., Hafen, E., Leevers, S., & Partridge, L. (2004). Long-lived Drosophila with over-expressed dFOXO in adult fat body. Science, 305(5682), 361. 10.1126/science.1098219

Gillespie, J., & Turelli, M. (1989). Genotype-environment interactions and the maintenance of polygenic variation. Genetics, 121(1), 129–138. 10.1093/genetics/121.1.129

Gomez-Lamarca, M., Falo-Sanjuan, J., Stojnic, R., Abdul Rehman, S., Muresan, L., Jones, M., Pillidge, Z., Cerda-Moya, G., Yuan, Z., Baloul, S., Valenti, P., Bystricky, K., Payre, F., O’Holleran, K., Kovall, R., & Bray, S. (2018). Activation of the Notch Signaling Pathway In Vivo Elicits Changes in CSL Nuclear Dynamics. Developmental Cell, 44(5), 611–623.e7. 10.1016/J.DEVCEL.2018.01.020/ATTACHMENT/98564200-F03C-488F-88AE-A3EEF35ED234/MMC3.MP4

Gulev, S., Thorne, P., Ahn, J., Dentener, F., Domingues, C., Gerland, S., Gong, D., Kaufman, D., Nnamchi, H., Quaas, J., Rivera, J., Sathyendranath, S., Smith, S., Trewin, B., von Schuckmann, K., & Vose, R. (2021). Changing State of the Climate System. En V. Masson-Delmotte, P. Zhai, A. Pirani, S. Connors, C. Péan, S. Berger, N. Caud, Y. Chen, L. Goldfarb, M. Gomis, M. Huang, K. Leitzell, E. Lonnoy, J. Matthews, T. Maycock, T. Waterfield, O. Yelekçi, R. Yu, & B. Zhou (Eds.), Climate Change 2021: The Physical Science Basis. Contribution of Working Group I to the Sixth Assessment Report of the Intergovernmental Panel on Climate Change (pp. 287-422). Cambridge University Press.

Hangartner, S., & Hoffmann, A. (2016). Evolutionary potential of multiple measures of upper thermal tolerance in Drosophila melanogaster. Functional Ecology, 30(3), 442–452. 10.1111/1365-2435.12499

Hartmann, D., Klein Tank, A., Rusticucci, M., Alexander, L., Brönnimann, S., Charabi, Y., Dentener, F., Dlugokencky, E., Easterling, D., Kaplan, A., Soden, B., Thorne, P., Wild, M., & Zhai, P. (2013). Observations: Atmosphere and surface. En T. Stocker, D. Qin, G. Plattner, M. Tignor, S. Allen, J. Boschung, A. Nauels, Y. Xia, V. Bex, & P. Midgley (Eds.), Climate Change 2013 the Physical Science Basis: Working Group I Contribution to the Fifth Assessment Report of the Intergovernmental Panel on Climate Change (pp. 159–254). Cambridge University Press. 10.1017/CBO9781107415324.008

Hoffmann, A., & Willi, Y. (2008). Detecting genetic responses to environmental change. Nature Reviews Genetics, 9(6), 421–432. 10.1038/nrg2339

Honjo, K., Mauthner, S., Wang, Y., Skene, J., & Tracey, W. (2016). Nociceptor-Enriched Genes Required for Normal Thermal Nociception. Cell Reports, 16(2), 295–303. 10.1016/J.CELREP.2016.06.003/ATTACHMENT/6B7A1D54-78AC-4973-A976-7EABB0439886/MMC2.XLSX

Huang, W., Campbell, T., Carbone, M. A., Jones, W. E., Unselt, D., Anholt, R., & Mackay, T. (2020). Context-dependent genetic architecture of Drosophila life span. PLoS Biology, 18(3), e3000645. 10.1371/journal.pbio.3000645

Huang, W., Massouras, A., Inoue, Y., Peiffer, J., Ràmia, M., Tarone, A., Turlapati, L., Zichner, T., Zhu, D., Lyman, R., Magwire, M., Blankenburg, K., Carbone, M. A., Chang, K., Ellis, L., Fernandez, S., Han, Y., Highnam, G., Hjelmen, C., … Mackay, T. (2014). Natural variation in genome architecture among 205 Drosophila melanogaster Genetic Reference Panel lines. Genome Research, 24(7), 1193–1208. 10.1101/gr.171546.113

Jørgensen, L. B., Malte, H., Ørsted, M., Klahn, N. A., & Overgaard, J. (2021). A unifying model to estimate thermal tolerance limits in ectotherms across static, dynamic and fluctuating exposures to thermal stress. Scientific Reports 2021 11:1, 11(1), 1-14. 10.1038/s41598-021-92004-6

Jørgensen, L. B., Malte, H., & Overgaard, J. (2019). How to assess Drosophila heat tolerance: Unifying static and dynamic tolerance assays to predict heat distribution limits. Functional Ecology, 33(4), 629–642. 10.1111/1365-2435.13279

Kapun, M., Barrón, M. G., Staubach, F., Obbard, D. J., Wiberg, R. A. W., Vieira, J., Goubert, C., Rota-Stabelli, O., Kankare, M., Bogaerts-Márquez, M., Haudry, A., Waidele, L., Kozeretska, I., Pasyukova, E. G., Loeschcke, V., Pascual, M., Vieira, C. P., Serga, S., Montchamp-Moreau, C., … González, J. (2020). Genomic Analysis of European Drosophila melanogaster Populations Reveals Longitudinal Structure, Continent-Wide Selection, and Previously Unknown DNA Viruses. Molecular Biology and Evolution, 37(9), 2661–2678. 10.1093/molbev/msaa120

Kellermann, V., Overgaard, J., Hoffmann, A., Fljøgaard, C., Svenning, J., & Loeschcke, V. (2012). Upper thermal limits of Drosophila are linked to species distributions and strongly constrained phylogenetically. Proceedings of the National Academy of Sciences of the United States of America, 109(40), 16228–16233. 10.1073/PNAS.1207553109/-/DCSUPPLEMENTAL

Kelly, M. (2019). Adaptation to climate change through genetic accommodation and assimilation of plastic phenotypes. Philosophical Transactions of the Royal Society B: Biological Sciences, 374(1768), 20180176. 10.1098/rstb.2018.0176

Kelly, M., Sanford, E., & Grosberg, R. (2012). Limited potential for adaptation to climate change in a broadly distributed marine crustacean. Proceedings of the Royal Society B: Biological Sciences, 279(1727), 349–356. 10.1098/RSPB.2011.0542

Kolaczkowski, B., Kern, A. D., Holloway, A. K., & Begun, D. J. (2011). Genomic Differentiation Between Temperate and Tropical Australian Populations of *Drosophila melanogaster*. Genetics, 187(1), 245–260. 10.1534/genetics.110.123059

Kuznetsova, A., Brockhoff, P. B., & Christensen, R. H. B. (2017). lmerTest Package: Tests in Linear Mixed Effects Models. Journal of Statistical Software, 82(13), 1–26. 10.18637/jss.v082.i13

Lansdell, S., Collins, T., Goodchild, J., & Millar, N. (2012). The Drosophila nicotinic acetylcholine receptor subunits Dα5 and Dα7 form functional homomeric and heteromeric ion channels. BMC Neuroscience, 13(1), 1–11. 10.1186/1471-2202-13-73/FIGURES/4

Lasne, C., Hangartner, S., Connallon, T., & Sgrò, C. (2018). Cross-sex genetic correlations and the evolution of sex-specific local adaptation: Insights from classical trait clines in Drosophila melanogaster. Evolution, 72(6), 1317–1327. 10.1111/evo.13494

Lecheta, M., Awde, D., O’Leary, T., Unfried, L., Jacobs, N., Whitlock, M., McCabe, E., Powers, B., Bora, K., Waters, J., Axen, H., Frietze, S., Lockwood, B., Teets, N., & Cahan, S. (2020). Integrating GWAS and Transcriptomics to Identify the Molecular Underpinnings of Thermal Stress Responses in Drosophila melanogaster. Frontiers in Genetics, 11, 658. 10.3389/fgene.2020.00658

Lenoir, J., Bertrand, R., Comte, L., Bourgeaud, L., Hattab, T., Murienne, J., & Grenouillet, G. (2020). Species better track climate warming in the oceans than on land. Nature Ecology & Evolution, 4(8), Article 8. 10.1038/s41559-020-1198-2

Lerman, D., & Feder, M. (2001). Laboratory selection at different temperatures modifies heat-shock transcription factor (HSF) activation in Drosophila melanogaster. Journal of Experimental Biology, 204(2), 315–323.

Li, A., Li, L., Zhang, Z., Li, S., Wang, W., Guo, X., & Zhang, G. (2021). Noncoding Variation and Transcriptional Plasticity Promote Thermal Adaptation in Oysters by Altering Energy Metabolism. Molecular Biology and Evolution, 38(11), 5144–5155. 10.1093/molbev/msab241

Li, Y. J., Chen, S. Y., Jorgensen, L. B., Overgaard, J., Renault, D., Colinet, H., & Ma, C. S. (2023). Interspecific differences in thermal tolerance landscape explain aphid community abundance under climate change. JOURNAL OF THERMAL BIOLOGY, 114. 10.1016/j.jtherbio.2023.103583

Lin, Y. J., Seroude, L., & Benzer, S. (1998). Extended life-span and stress resistance in the Drosophila mutant methuselah. Science, 282(5390), 943–946. 10.1126/science.282.5390.943

Loeschcke, V., Kristensen, T., & Norry, F. (2011). Consistent effects of a major QTL for thermal resistance in field-released Drosophila melanogaster. Journal of Insect Physiology, 57(9), 1227–1231. 10.1016/j.jinsphys.2011.05.013

Logan, M., & Cox, C. (2020). Genetic Constraints, Transcriptome Plasticity, and the Evolutionary Response to Climate Change. Frontiers in Genetics, 11, 1088. 10.3389/FGENE.2020.538226/BIBTEX

Lynch, M., & Walsh, B. (1998). *Genetics and analysis of quantitative traits*. Sunauer Associates, Inc.

Machado, H. E., Bergland, A. O., O’Brien, K. R., Behrman, E. L., Schmidt, P. S., & Petrov, D. A. (2016). Comparative population genomics of latitudinal variation in *Drosophila simulans* and *Drosophila melanogaster*. Molecular Ecology, 25(3), 723–740. 10.1111/mec.13446

MacKay, T., Richards, S., Stone, E., Barbadilla, A., Ayroles, J., Zhu, D., Casillas, S., Han, Y., Magwire, M., Cridland, J., Richardson, M., Anholt, R., Barrón, M., Bess, C., Blankenburg, K., Carbone, M., Castellano, D., Chaboub, L., Duncan, L., … Gibbs, R. (2012). The Drosophila melanogaster Genetic Reference Panel. Nature, 482(7384), 173–178. 10.1038/nature10811

McColl, G., & McKechnie, S. (1999). The Drosophila heat shock hsr-omega gene: An allele frequency cline detected by quantitative PCR. Molecular Biology and Evolution, 16(11), 1568–1574. 10.1093/oxfordjournals.molbev.a026069

Meehl, G., & Tebaldi, C. (2004). More intense, more frequent, and longer lasting heat waves in the 21st century. Science, 305(5686), 994–997. 10.1126/science.1098704

Mesas, A., Jaramillo, A., & Castañeda, L. (2021). Experimental evolution on heat tolerance and thermal performance curves under contrasting thermal selection in Drosophila subobscura. Journal of Evolutionary Biology, 34(5), 767–778. 10.1111/JEB.13777

Miech, C., Pauer, H., He, X., & Schwarz, T. (2008). Presynaptic local signaling by a canonical wingless pathway regulates development of the Drosophila neuromuscular junction. The Journal of neuroscience : the official journal of the Society for Neuroscience, 28(43), 10875–10884. 10.1523/JNEUROSCI.0164-08.2008

Mitchell, K. A., & Hoffmann, A. A. (2010). Thermal ramping rate influences evolutionary potential and species differences for upper thermal limits in Drosophila. Functional Ecology, 24(3), 694–700. 10.1111/J.1365-2435.2009.01666.X

Morgan, T., & Mackay, T. (2006). Quantitative trait loci for thermotolerance phenotypes in Drosophila melanogaster. Heredity, 96(3), 232–242. 10.1038/sj.hdy.6800786

Norry, F., Gomez, F., & Loeschcke, V. (2007). Knockdown resistance to heat stress and slow recovery from chill coma are genetically associated in a quantitative trait locus region of chromosome 2 in *Drosophila melanogaster*. Molecular Ecology, 16(15), 3274–3284. 10.1111/j.1365-294X.2007.03335.x

Norry, F., Larsen, P., Liu, Y., & Loeschcke, V. (2009). Combined expression patterns of QTL-linked candidate genes best predict thermotolerance in Drosophila melanogaster. Journal of Insect Physiology, 55(11), 1050–1057. 10.1016/j.jinsphys.2009.07.009

Ørsted, M., Hoffmann, A. A., Rohde, P. D., Sørensen, P., & Kristensen, T. N. (2019). Strong impact of thermal environment on the quantitative genetic basis of a key stress tolerance trait. Heredity, 122(3), 315. 10.1038/S41437-018-0117-7

Ørsted, M., Jørgensen, L. B., & Overgaard, J. (2022). Finding the right thermal limit: A framework to reconcile ecological, physiological and methodological aspects of CTmax in ectotherms. Journal of Experimental Biology, 225(19). 10.1242/JEB.244514/277015

Overgaard, J., Hoffmann, A., & Kristensen, T. (2011). Assessing population and environmental effects on thermal resistance in Drosophila melanogaster using ecologically relevant assays. Journal of Thermal Biology, 36(7), 409–416. 10.1016/j.jtherbio.2011.07.005

Pacifici, M., Foden, W., Visconti, P., Watson, J., Butchart, S., Kovacs, K. M., Scheffers, B., Hole, D., Martin, T., Akçakaya, H. R., Corlett, R., Huntley, B., Bickford, D., Carr, J., Hoffmann, A., Midgley, G., Pearce-Kelly, P., Pearson, R., Williams, S., … Rondinini, C. (2015). Assessing species vulnerability to climate change. Nature Climate Change, 5(3), 215–225. 10.1038/nclimate2448

Pennell, T., De Haas, F., Morrow, E., & Van Doorn, G. (2016). Contrasting effects of intralocus sexual conflict on sexually antagonistic coevolution. Proceedings of the National Academy of Sciences of the United States of America, 113(8), E978–E986. 10.1073/pnas.1514328113

Pérez-Gálvez, F. R., Zhou, S., Wilson, A. C., Cornwell, C. L., Awde, D. N., & Teets, N. M. (2023). Scoring thermal limits in small insects using open-source, computer-asssited motion detection. Journal of Experimental Biology, 226(22): jeb246548. 10.1242/jeb.246548

Rezende, E., Bozinovic, F., Szilágyi, A., & Santos, M. (2020). Predicting temperature mortality and selection in natural Drosophila populations. Science, 369(6508), 1242–1245. 10.1126/SCIENCE.ABA9287

Rezende, E., Castañeda, L., & Santos, M. (2014). Tolerance landscapes in thermal ecology. Functional Ecology, 28(4), 799–809. 10.1111/1365-2435.12268

Rezende, E., Tejedo, M., & Santos, M. (2011). Estimating the adaptive potential of critical thermal limits: Methodological problems and evolutionary implications. Functional Ecology, 25(1), 111–121. 10.1111/j.1365-2435.2010.01778.x

Roff, D. (1997). Evolutionary Quantitative Genetics.

Rolandi, C., Lighton, J., de la Vega, G., Schilman, P., & Mensch, J. (2018). Genetic variation for tolerance to high temperatures in a population of *Drosophila melanogaster*. Ecology and Evolution, 8(21), 10374–10383. 10.1002/ece3.4409

Rudman, S. M., Greenblum, S. I., Rajpurohit, S., Betancourt, N. J., Hanna, J., Tilk, S., Yokoyama, T., Petrov, D. A., & Schmidt, P. (2022). Direct observation of adaptive tracking on ecological time scales in *Drosophila*. Science, 375(6586), eabj7484. 10.1126/science.abj7484

Ruzicka, F., Dutoit, L., Czuppon, P., Jordan, C., Li, X., Olito, C., Runemark, A., Svensson, E., Yazdi, H. P., & Connallon, T. (2020). The search for sexually antagonistic genes: Practical insights from studies of local adaptation and statistical genomics. Evolution Letters, 4(5), 398–415. 10.1002/evl3.192

Saltz, J. B., Bell, A. M., Flint, J., Gomulkiewicz, R., Hughes, K. A., & Keagy, J. (2018). Why does the magnitude of genotype-by-environment interaction vary? Ecology and Evolution, 8(12), 6342–6353. 10.1002/ECE3.4128

Sambucetti, P., Scannapieco, A., & Norry, F. (2010). Direct and correlated responses to artificial selection for high and low knockdown resistance to high temperature in Drosophila buzzatii. Journal of Thermal Biology, 35(5), 232–238. 10.1016/J.JTHERBIO.2010.05.006

Santos, M., Castañeda, L., & Rezende, E. (2012). Keeping pace with climate change: What is wrong with the evolutionary potential of upper thermal limits? Ecology and Evolution, 2(11), 2866–2880. 10.1002/ece3.385

Sgrò, C., Overgaard, J., Kristensen, T., Mitchell, K., Cockerell, F., & Hoffmann, A. (2010). A comprehensive assessment of geographic variation in heat tolerance and hardening capacity in populations of Drosophila melanogaster from Eastern Australia. Journal of Evolutionary Biology, 23(11), 2484–2493. 10.1111/j.1420-9101.2010.02110.x

Terblanche, J., Deere, J., Clusella-Trullas, S., Janion, C., & Chown, S. (2007). Critical thermal limits depend on methodological context. Proceedings of the Royal Society B: Biological Sciences, 274(1628), 2935–2942. 10.1098/rspb.2007.0985

Thomas, C., Cameron, A., Green, R., Bakkenes, M., Beaumont, L., Collingham, Y., Erasmus, B., Ferreira De Siqueira, M., Grainger, A., Hannah, L., Hughes, L., Huntley, B., Van Jaarsveld, A., Midgley, G., Miles, L., Ortega-Huerta, M., Peterson, T., Phillips, O., & Williams, S. (2004). Extinction risk from climate change. Nature, 427(6970), 145–148. 10.1038/nature02121

Timpson, N. J., Greenwood, C. M. T., Soranzo, N., Lawson, D. J., & Richards, J. B. (2017). Genetic architecture: The shape of the genetic contribution to human traits and disease. Nature Reviews Genetics 2017 19:2, 19(2), 110-124. 10.1038/nrg.2017.101

Turner, S. (2014). qqman: An R package for visualizing GWAS results using Q-Q and manhattan plots. bioRxiv, 005165. 10.1101/005165

Van Heerwaarden, B., Malmberg, M., & Sgrò, C. (2016). Increases in the evolutionary potential of upper thermal limits under warmer temperatures in two rainforest Drosophila species. Evolution, 70(2), 456–464. 10.1111/EVO.12843

Weber, A., Khan, G., Magwire, M., Tabor, C., Mackay, T., & Anholt, R. (2012). Genome-wide association analysis of oxidative stress resistance in Drosophila melanogaster. PloS one, 7(4). 10.1371/JOURNAL.PONE.0034745

Zhao, L., Wit, J., Svetec, N., & Begun, D. J. (2015). Parallel Gene Expression Differences between Low and High Latitude Populations of Drosophila melanogaster and D. simulans. PLOS Genetics, 11(5), e1005184. 10.1371/journal.pgen.1005184

